# Coherence Shifts in Attribute Evaluations

**DOI:** 10.1101/2020.08.19.258046

**Authors:** Douglas G. Lee, Keith J. Holyoak

## Abstract

In five experiments, people repeatedly judged individual options with respect to both overall value and attribute values. When required to choose between two snacks, each differing in two attributes (pleasure and nutrition), people’s assessments of value shifted from pre- to post-choice in the direction that spread the alternatives further apart so as to favor the winner, thereby increasing confidence in the choice. This shift was observed not only for ratings of overall value, but also for each of the two individual attributes. The magnitude of the coherence shift increased with choice difficulty as measured by the difference in initial ratings of overall value for the two options, as well as with a measure of attribute disparity (the degree to which individual attributes “disagree” with one another as to which option is superior). In Experiments 2-5, tasks other than explicit choice generated the same qualitative pattern of value changes, confidence, and response time. These findings support the hypothesis that active consideration of options, whether or not explicitly related to value, automatically refines the mental value representations for the options, which in turn allows them to be more precisely distinguished when later included in a value-based choice set.

## Introduction

The traditional view of multi-attribute decision making assumes that when people choose between options, they compare them with respect to a set of decision-relevant attributes. According to one theoretical perspective, the attributes of each option are assessed individually, weighted according to their importance, and summed together to map onto an estimate of overall value for each option (e.g., Tversky & Simonson, 1993; Rieskamp, Busemeyer, & Mellers, 2006). Alternatively, attributes may be assessed and compared separately across options, with differences at the level of individual attributes being used to choose the preferred option (e.g., Tversky, 1969; Hunt, Dolan, & Behrens, 2014). In either case, attribute values constitute pre-existing and stable elements that serve as the basic inputs to the decision process.

An alternative perspective emphasizes the constructive nature of preference (e.g., Ariely, Loewenstein, & Prelec, 2006; DeKay, Stone, & Miller, 2011; Holyoak & Simon, 1999; Russo, Medvec, & Meloy, 1996; Payne, Bettman, & Schkade, 1999; Svenson, 1992; also see Lichtenstein & Slovic, 2006; Warren, McGraw, & Van Boven, 2011). Under this view, subjective assessments of attribute values and attribute importance can shift during the decision process so as to cohere with the emerging decision. These shifts will spread the perceived values of the options, increasing the advantage of the eventual winner relative to its competitor(s) and thereby increasing confidence in the decision. Such spreading of alternatives (SoA), or coherence shifts, have been shown to impact decisions about such everyday matters as choosing a restaurant or an apartment to rent (Russo et al., 1996; Simon, Krawzcyk, & Holyoak, 2004; Simon, Krawczyk, Bleicher, & Holyoak, 2008), adjudicating legal disputes (Holyoak & Simon, 1999; Carlson & Russo, 2001; Simon, 2012), and evaluating complex issues with moral implications (Spellman, Ullman, & Holyoak, 1993; Holyoak & Powell, 2015; Simon, Stenstrom, & Read, 2015). Cognitive evaluations and emotional reactions can jointly undergo coherence shifts during decision making (Simon, Stenstrom, & Read, 2015). In contrast to classical cognitive dissonance theory (Festinger, 1957; see Harmon-Jones & Harmon-Jones, 2007, for a review), which claims that value shifts are post-decisional, the construction of preference emphasizes that coherence shifts play a critical role in the pre-decisional dynamics that drive the selection of an option (Simon & Holyoak, 2002). A number of studies have shown that coherence shifts are observed prior to making a firm decision or public commitment (Holyoak & Simon, 1999; Russo et al., 1996; Russo, Carlson, Meloy, & Yong, 2008; Russo & Chaxel, 2010; Simon et al., 2004; Simon, Pham, Le, & Holyoak, 2001).

Although the impact of coherence shifts on decision making has been firmly established, less is known about the intra-decisional dynamics that drive attribute reevaluation. The way in which people process information during decision making may be subject to metacognitive control (e.g., Chaxel, 2015), which may lead to *rational inattention* to different attributes in different choice contexts (Sims, 2003; Caplin & Dean, 2015). The magnitude of shifts in overall evaluations of options tends to increase with the difficulty of the decision (Svenson, 1992), thereby increasing confidence in the choice (Simon, Snow & Read, 2004; Simon & Spiller, 2016; Lee & Daunizeau, 2020a). However, previous studies relating choice difficulty to the magnitude of coherence shifts have not assessed potential changes in the perceived values of those individual attributes that are hypothesized to determine overall value. In the present paper, we report a series of experiments that used a multi-attribute choice paradigm to examine coherence shifts in both overall evaluations and in perceived values of individual attributes. We expected that more difficult choices—those in which two options are initially relatively close in overall value—will trigger larger coherence shifts both for overall value and for individual attributes.

In addition, we examined the impact of a different factor potentially related to choice difficulty. We hypothesized that decision dynamics may also be influenced by differences in attribute composition. Consider the two hypothetical choices situations depicted in Figure 1. Both choices involve two options based on two relevant and equally-weighted attributes, A1 and A2. The first choice is between option 1 with attribute values of 90 for A1 and 30 for A2, versus option 2 with values 20 and 80. The second choice is between option 3 with values 65 and 55, versus option 4 with values 45 and 55. Under the assumption of equal importance weights for A1 and A2, the two choices are identical at the level of overall value (60 for option 1 and option 3, 50 for option 2 and option 4). Thus option 1 is favored for the first choice, and option 3 for the second choice, by equal amounts. But notice that the disparity between the attributes is much greater for the first choice (option 1 versus option 2) than for the second (option 3 versus option 4).

**Figure 1.**
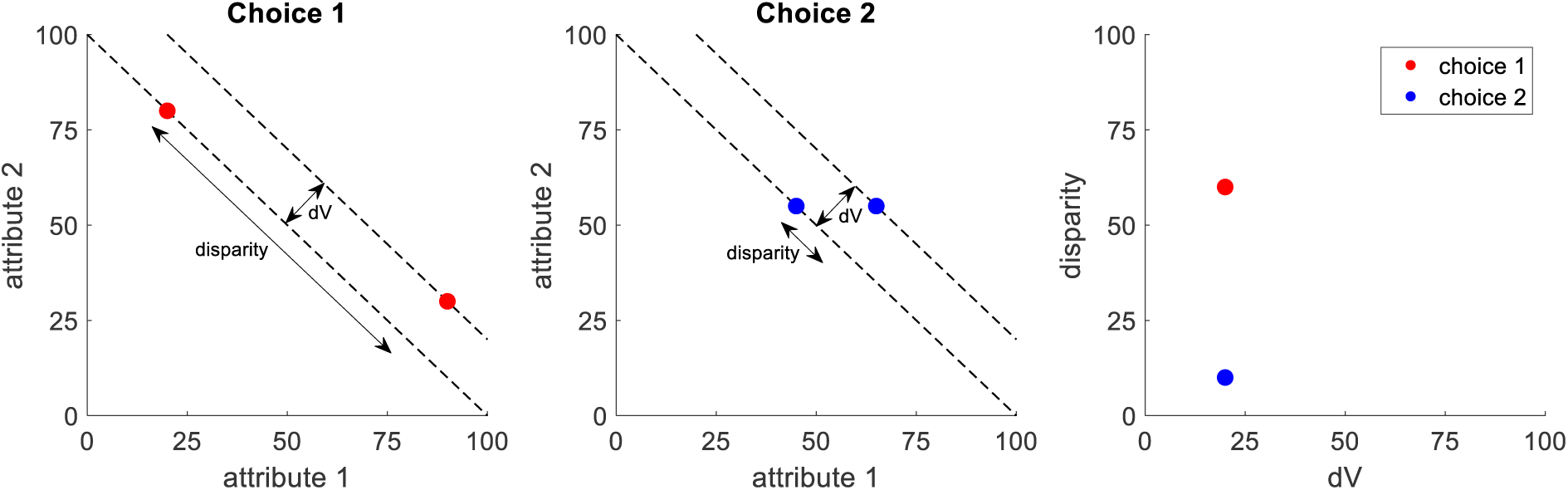
A schematic illustration of orthogonal components of choice difficulty: dV and disparity. The left plot illustrates a “high disparity” choice and the middle plot illustrates a “low disparity” choice. The red and blue dots represent the alternative options for each choice, each plotted according to its measurements on two attribute dimensions. The example assumes equal importance weights for each attribute, so the iso-value curves are represented by parallel lines with slope −1. The difference in overall value of the options, dV, is the distance between the iso-value curves on which the options lie. Disparity is the distance between the options in the dimension orthogonal to overall value (see Equation 1 for a mathematical formulation). The right plot shows the location of each choice pair in the transformed dV-disparity space.

We propose that the choice with higher disparity will result in larger coherence shifts and thus greater confidence. When choice options are disparate, one option dominates along one attribute dimension, whereas the other dominates along the other dimension. It has been argued that the more dominant is an attribute of a particular option (relative to another option or the average across all other options), the more attention it will receive during the deliberation leading to a decision (Bordalo, Gennaioli, & Shleifer, 2020, 2013). Such increased attention would be expected to enhance the processes that produce spreading of alternatives. An increased coherence shift for high disparity choices would in turn increase choice confidence and decrease response time (RT), as the choice effectively becomes easier (cf. Lee & Daunizeau, 2020a).

In addition to addressing the above issues, the present study attempted to identify the core component of the decision-making mechanisms that generates coherence shifts. Coherence shifts evidenced by SoA have typically been assessed in a choice context (hence SoA is commonly referred to as *choice-induced preference change* (CIPC); Izuma et al., 2010; Voigt, Murawski, Speer, & Bode, 2019). There is evidence, however, that coherence shifts can be produced by meaningful processing of options without introducing an overt choice task (DeKay 2015; DeKay, Stone, & Sorenson, 2012; Janis & Mann, 1977; Montgomery & Willen, 1999; Russo et al., 1996, 2008; Simon et al., 2001; Svenson, 1992; for reviews see Brownstein, 2003; Russo, 2015). Moreover, it has been suggested that CIPC may actually be an illusion based on a statistical artifact of the rating➔choice➔rating (RCR) task design. A study by Chen and Risen (2010) showed that choices between items that are (subjectively) rated as being very close in value will lead to apparent CIPC (on average), even if there is no true cognitive basis for the effect. If the latent mental representations of the option values are assumed to be imprecise (i.e., they have a variance around their expected value), sometimes the initial rating for the to-be chosen item will be drawn from the low tail of its value distribution, and the initial rating for the to-be rejected item will be drawn from the high tail of its value distribution. The final ratings, however, would most likely be closer to the “true” values (i.e., the means of the respective value distributions), due to the laws of probability. In this case, the observed CIPC would have been caused by a statistical artifact of the sampling procedure, not cognitive reflection. Chen and Risen introduced a new experimental condition to control for this statistical explanation, rating➔rating➔choice (RRC). In accord with their statistical hypothesis, they found a significant level of CIPC even in this control condition, thus proving their point. However, a number of subsequent studies have shown that CIPC in the standard RCR condition is significantly higher than in the RRC condition, suggesting that choosing between options does indeed cause CICP beyond what can be explained by statistical considerations (Lee & Daunizeau, 2020a; Chammat et al., 2017; Salti et al., 2014; Coppin et al., 2014; Sharot et al., 2012).

Some degree of CIPC can thus occur without any cognitive changes in value assessments (Chen & Risen, 2010). It has been suggested that deliberation might induce CIPC as option value representations are refined (Lee & Daunizeau, 2020b). However, there has as yet been no investigation of the extent to which different types or different degrees of refinement could take place during tasks other than explicit choice deliberation. We hypothesize that the refinement of value representations is a continuous rather than binary phenomenon, and that different tasks will elicit degrees of coherence shifts that fall along a spectrum. The low end of this spectrum will occur when there is no task other than repeated ratings (e.g., the RRC control); the high end of the spectrum should result when there is explicit deliberation about which option to choose (e.g., the standard RCR). Other tasks may result in intermediate degrees of coherence shifts, in proportion to the amount of value-relevant information processing that takes place. We propose that the core component of decision making that triggers coherence shifts is active consideration of the options, whether or not such consideration is explicitly related to value. Clearly, the more the task at hand resembles a choice (e.g., assessing similarity of the values of two options; see Experiment 2), the more value-related refinement should occur. Tasks that involve comparing options but not explicitly on value (e.g., generic similarity judgments; see Experiment 3) should lead to a moderate level of value-related refinement. Tasks that neither focus on value nor require comparison of options (see Experiments 4-5) should lead to lower (but still detectable) degrees of refinement. Such a pattern would be consistent with previous work showing that merely repeating isolated evaluations of options causes the evaluations to become more precise (Lee & Coricelli, 2020).

We conducted a series of behavioral experiments in which people judged individual options with respect to both overall value and attribute values, both prior to and after making a choice (Experiment 1) or performing some other task based on the same set of stimuli (Experiments 2-5). In the main experiment (Experiment 1), we assessed how the relationship between the options (in terms of both difference in overall value and in attribute disparity) impacted eventual choice, choice confidence, RT, and shifts in overall value and in attribute values. In the auxiliary experiments (Experiments 2-5), we assessed how the degree of coherence shifts varied as a function of (assumed) value-relevant information processing.

## Experiment 1

Experiment 1 examined shifts in overall option values and in attribute values triggered by the choice between two snacks with varying initial values on two attributes, pleasure and nutrition.

### Method

#### Participants

A total of 58 people participated in Experiment 1 (32 female; age: mean = 39 years, SD = 8, range 25-50). (The data for one participant were corrupt, so we excluded this participant from analyses.) This sample size was chosen to be comparable to that used in previous studies based on a similar paradigm. All participants were recruited using Amazon Mechanical Turk (MTurk) (https://mturk.com). All were classified as “masters” by Mturk. They were residents of the United States or Canada, and all were native English speakers. Each participant received a payment of $7.50 as compensation for approximately one hour of time. Our experiments involved de-identified participant data, and protocols were approved by the Institutional Review Board of the University of California, Los Angeles. All participants gave informed consent prior to commencing the experiments.

#### Materials

The experiments were constructed using the online experimental platform Gorilla (gorilla.sc). The experimental stimuli were drawn from a set of 200 digital images used in a previous study (Lee & Coricelli, 2020), each representing a distinct snack food item. For Experiment 1, we used a subset of 100 stimuli, identical for all participants. We determined this subset by selecting the 100 items for which ratings from the previous study varied the least (within each item) across participants. Figure 2 shows examples of images of snacks, indicating their initial values on the two attributes of pleasure and nutrition (averaged across participants).

**Figure 2.**
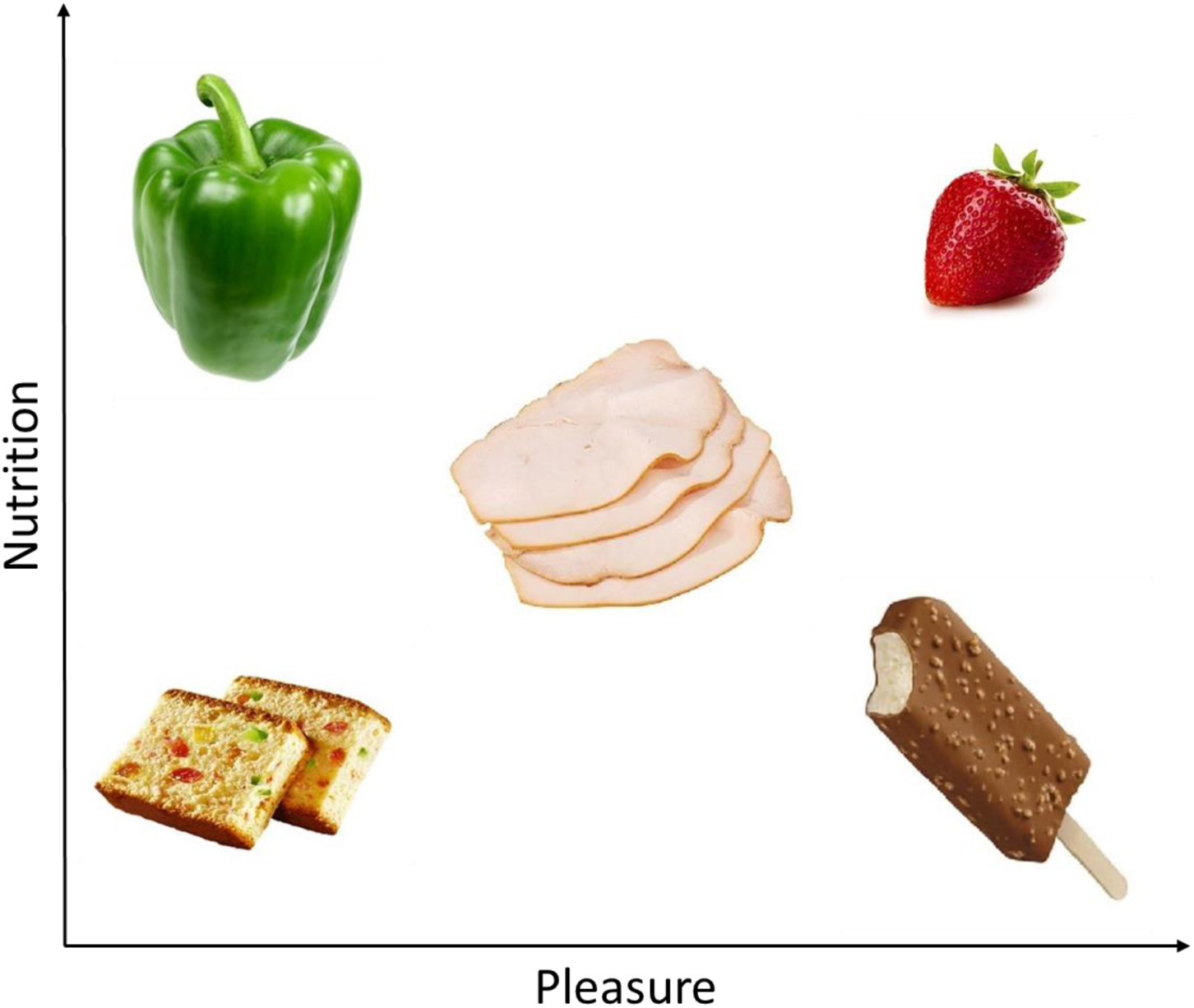
An example of snack foods included in the stimulus set, plotted according to their respective ratings along the dimensions of pleasure and nutrition (averaged across participants). From lower left to upper right: fruitcake rated low on both pleasure and nutrition; bell pepper rated low on pleasure but high on nutrition; sliced turkey rated medium on both pleasure and nutrition; ice cream bar rated high on pleasure but low on nutrition; strawberry rated high on both pleasure and nutrition.

#### Design and Procedure

The experiment consisted of a pre-exposure phase, followed by initial value ratings, choice, and final ratings. No time limits were imposed for any of the constituent tasks or for the overall experiment.

In the pre-exposure phase, participants simply observed as all individual items were displayed in a random sequence for 750 ms each. The purpose of the pre-exposure phase was to familiarize participants with the full set of items that they would later evaluate, allowing them to form an impression of the range of subjective values across the item set.

The initial value ratings comprised three sets of ratings: for overall value of the snack, and for the value of each of two individual attributes, nutrition and pleasure. In each rating task, all stimuli were displayed on the screen, one at a time, in a sequence randomized across participants. At the onset of each trial, a fixation cross appeared at the center of the screen for 750 ms. Next, an image of a single food item appeared at the center of the screen. For the rating of overall value, participants responded to the question, “How much would you like this as a daily snack?” using a horizontal slider scale. This question was intended to motivate participants to think carefully while assessing the overall subjective quality of each option, as the choice was to determine long-term daily consumption, rather than a “one off” snack. The leftmost end of the scale was labeled “Not at all,” and the rightmost end was labeled “Very much!” The scale appeared to participants to be continuous, and the data was captured in increments of 1 (ranging from 1 to 100). Participants could revise their rating as many times as they liked before finalizing it. Participants clicked the “enter” button to finalize their value rating response and proceed to the next screen. The next trial then began.

The overall value ratings were followed by attribute ratings, which were performed separately for nutrition and pleasure, with the order counterbalanced across participants. One attribute was rated for every alternative (in one task section), then the other attribute was rated for every alternative (in the next task section). In each attribute rating task, all stimuli were displayed on the screen, one at a time, in a random sequence (randomized across participants and across sections for each participant). The format of this task was identical to that of the overall value rating task, except now participants responded to the question, “How nutritious do you consider this item to be?” or “How pleasurable do you consider this item to be?” For both attribute ratings, the leftmost end of the slider scale was labeled “Very low!” and the rightmost end was labeled “Very high!”.

The choice task was then administered. For this task, 50 pairs of stimuli were displayed on the screen, one pair at a time, in a sequence randomized across participants. The pairings of items for each choice trial were created so as make the choices relatively difficult, as assessed by small differences in value ratings between the two items in a choice pair in a previous study (Lee & Coricelli, 2020). Each individual item occurred in a single choice pair. At the onset of each trial, a fixation cross appeared at the center of the screen for 750 ms. Next, a pair of images of food items appeared on the screen, one left and one right of center. Participants responded to the question, “Which would you prefer as a daily snack?” by clicking on the image of their preferred item. Participants then responded to the question, “How sure are you about your choice?” using a horizontal slider scale. The leftmost end of the scale was labeled “Not at all!” and the rightmost end was labeled “Absolutely!” Participants could revise their confidence report as many times as they liked before finalizing it. Participants clicked the “enter” button to finalize their confidence report and proceed to the next screen.

Finally, participants made final ratings of overall value and attribute values, exactly as in the initial ratings, except with stimuli presented in new random orders. Note that this procedure (randomizing order of individual items) serves to separate the ratings from the prior choice context. Prior to completing these final ratings, participants were instructed not to try to remember their earlier ratings, but rather to simply rate the stimuli as they now evaluated them.

### Results and Discussion

Due to the lack of experimental control in online experiments, we anticipated that not all participants would perform the tasks properly. In particular, we received feedback from some participants that the experiment was rather tedious, due to the repetitive nature required by its design. We therefore performed checks to assess whether participants might have given sloppy responses in the later sections of the experiment. Specifically, we calculated the Spearman correlation between first and last ratings, within participant. We also used generalized linear model (GLM) logistic regression of value ratings on choice to calculate the slope coefficient for each participant (separately using first and last ratings). Using these two measures, we deemed participants with scores outside a cutoff (median +/− 3 * median average deviation) to be an outlier, and removed them from our analyses. This procedure resulted in the removal of 8 participants for Experiment 1. All analyses were therefore based on 50 remaining participants.

Unless stated otherwise, all statistical effect sizes reported below were first calculated at the individual level and then tested for significance at the population level. All reported *p*-values represent the probability of non-zero effect sizes, based on standard two-sided *t*-tests. (See the Supplementary Material for tables containing detailed statistical summaries for each of the effects reported below.) For every GLM regression analysis reported below, all variables were first converted to *z*-scores (within participant) before being entered into the models. To assist with both readability and interpretation, we coded all variables such that option 1 (for each choice) refers to the option with the higher overall value rating in the first phase. We thus define dV (value difference) as the difference in overall value ratings (option 1 minus option 2). We define dP (pleasure difference) and dN (nutrition difference) in an analogous manner.

#### Coherence Shifts in Overall Value

Our primary analysis focus is on coherence shifts, which result in spreading of alternatives (SoA) from initial to final ratings. The choice defines the winning option, and SoA is defined in terms of changes that relatively favor the winner. SoA can be defined for both overall value ratings and for ratings of individual attribute values. For overall value, SoA is defined as the change in overall value for the chosen option from initial to final rating minus the change in overall value for the unchosen option. In accord with previous work showing coherence shifts over the course of decision making, we observed a reliable SoA in overall value across all choice trials (cross-participant mean of within-participant median SoA = 2.82, *p* < .001), which is comparable to the magnitude observed in previous studies (Lee & Daunizeau, 2020a; Voigt et al., 2019; Izuma et al., 2010). (See the Supplementary Material for more details on SoA.)

We then assessed the relationship between choice difficulty measured by the difference in initial overall value ratings between the options, *dV_1_*, where the subscript distinguishes the initial ratings (1) from the final ratings (2). As in previous studies (Lee & Coricelli, 2020; Lee & Daunizeau, 2020a, 2020b), dV_1_ had a reliable negative relationship with SoA and RT, and a reliable positive relationship with choice consistency and choice confidence. We used GLM to regress dV_1_ on overall SoA, on RT, on consistency, and on confidence, separately. The beta weights for dV_1_ on each dependent variable were significant and in the predicted direction: mean beta for SoA = −0.247 (*p* < .001); mean beta for RT = −0.218 (*p* < .001); mean beta for consistency = 1.634 (*p* < .001); mean beta for confidence = 0.354 (*p* < .001).

Lee and Daunizeau (2020a) proposed that decision makers refine their value estimates and certainty for the choice options during deliberation, prior to committing to the choice. In support of this hypothesis, these investigators showed that the impact of dV on RT, on consistency, and on confidence is larger when dV is calculated using post-choice ratings (i.e., dV_2_) rather than pre-choice ratings. This basic finding (also observed by Simon et al., 2004; Simon & Spiller, 2016) was replicated in our Experiment 1. The mean GLM beta weight for dV_2_ as a predictor of RT was −0.2478 (*p* < .001). The magnitude of this beta weight was greater than that obtained using dV_1_ (*p* = .037, one-sided t-test). The mean GLM beta weight for dV_2_ as a predictor of consistency was 2.7106 (*p* < .001). The magnitude of this beta weight was reliably greater than that obtained using dV_1_ (*p* < .001, one-sided t-test). The mean GLM beta weight for dV_2_ as a predictor of confidence was 0.3955 (*p* < .001). The magnitude of this beta weight was reliably greater than that obtained using dV_1_ (*p* = .015, one-sided t-test).

Lee and Daunizeau (2020a) also found that SoA positively influences choice confidence, as would be predicted given that the impact of a coherence shift is to effectively make the choice easier prior to entering a response. In brief, a high value of dV_1_ will make the choice an easy one, and confidence will therefore be high. A low dV_1_ will make the choice difficult, thereby encouraging deliberation before responding. Deliberation will on average result in a large SoA (which is essentially an increment in dV_1_), and therefore will lead to higher confidence. When we included both dV_1_ and SoA in a GLM regression model to predict confidence, the cross-participant mean beta weight for dV_1_ was 0.397 (*p* < .001) and for SoA was 0.168 (*p* < .001). We also checked for a similar effect of SoA on RT. When we included both dV_1_ and SoA in a GLM regression model to predict RT, the cross-participant mean beta weight for dV_1_ was −0.241 (*p* < .001) and for SoA was −0.062 (*p* = .013).

Finally, we tested our predictions that attribute disparity would correlate negatively with both SoA and choice confidence, and positively with RT. To explicitly define our disparity variable, we transformed the space representing choice options from attribute space (i.e., a two-dimensional space composed of pleasure and nutrition axes) to a new space in which one dimension is dV_1_ and the other is a value of disparity. Each point in the original attribute space (representing an individual item) necessarily resides on a specific iso-value line (i.e., an imaginary line that connects all points of equal overall value), the slope of which is determined by the (participant-specific) marginal rate of substitution (MRS) of the attributes. We calculated the MRS for each participant as –b_P_/b_N_, where b_P_ and b_N_ are the beta weights from the regression of pleasure and nutrition on overall value. The difference in the overall value of the two options being compared is thus the distance between the two iso-value lines on which the options reside (measured in the direction orthogonal to the lines). The disparity measure that we seek is the distance between the two options in the direction parallel to the iso-value lines (see Figure 3 for an illustrative example). This is simply the distance between the scalar projections of the two points (i.e., the location of the options in attribute space) onto the iso-value vector ([-b_P_ b_N_]’):

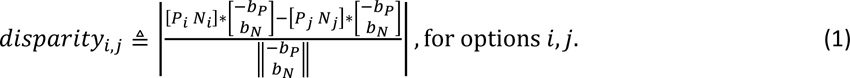

**Figure 3.**
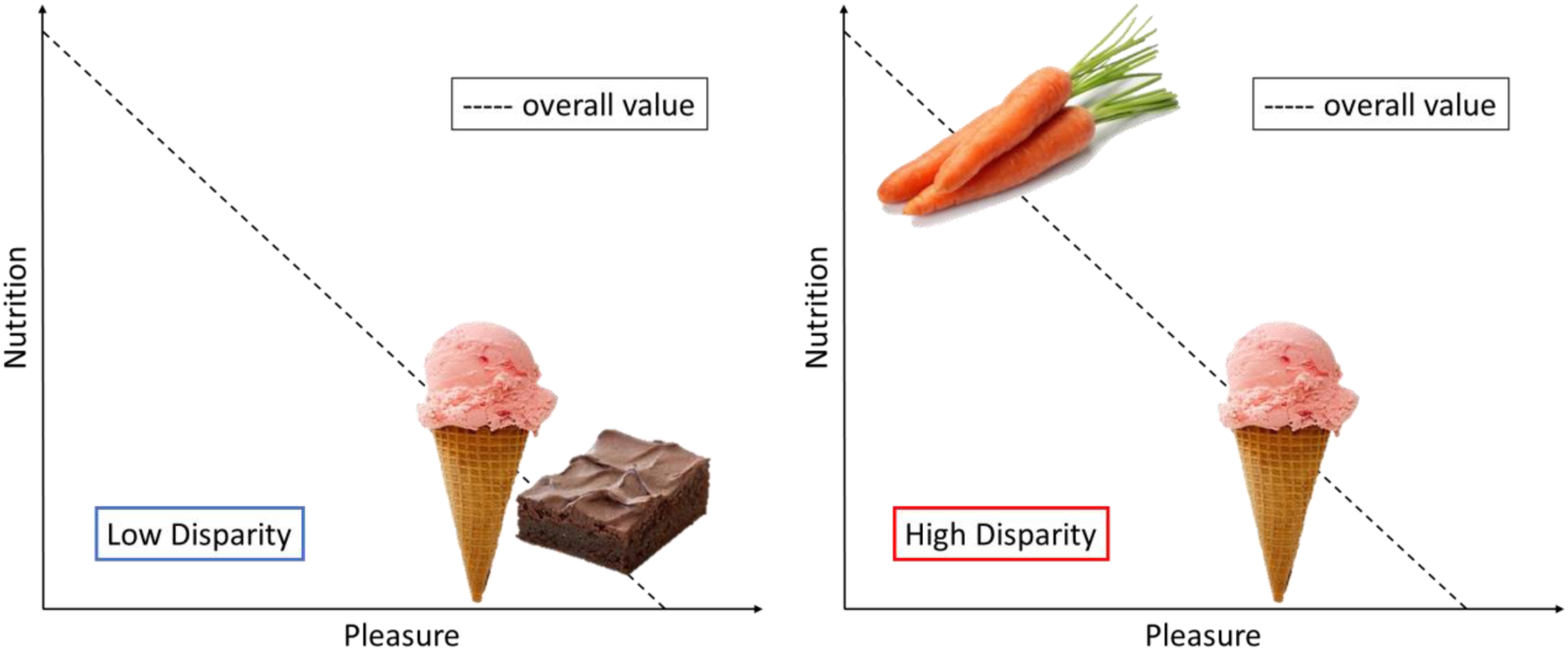
An illustration of two choice sets for snack foods, one low disparity (left plot), one high disparity (right plot). As shown by the dashed iso-value lines, all of the available snacks are of comparable overall value (and thus each choice pair is of comparable low dV). However, the two choice pairs are of very different disparity. In the low disparity pair (left), both options score high on pleasure and low on nutrition. In the high disparity pair (right), one option scores high on pleasure but low on nutrition, while the other option scores low on pleasure but high on nutrition.

Using this formal definition of disparity, we performed GLM regressions at the participant level of dV_1_ and disparity_1_ (where the subscript indicates disparity based on initial ratings; for simplicity we henceforth drop the subscripts for the two options) on SoA, RT, choice consistency, and choice confidence. Across participants, all beta weights were significant and in the predicted direction, though the impact of disparity on RT was not significant (mean dV_1_ beta for SoA = - 0.344, *p* < .001; mean disparity_1_ beta for SoA = 0.184, *p* < .001; mean dV_1_ beta for RT = −0.195, *p* < .001; mean disparity_1_ beta for RT = −0.037, *p* =.112 mean dV_1_ beta for consistency = 1.558, *p* < .001; mean disparity_1_ beta for consistency = 0.249, *p* =.022; mean dV_1_ beta for confidence = 0.313, *p* < .001; mean disparity_1_ beta for confidence = 0.071, *p* =.026; see Figure 4).

**Figure 4.**
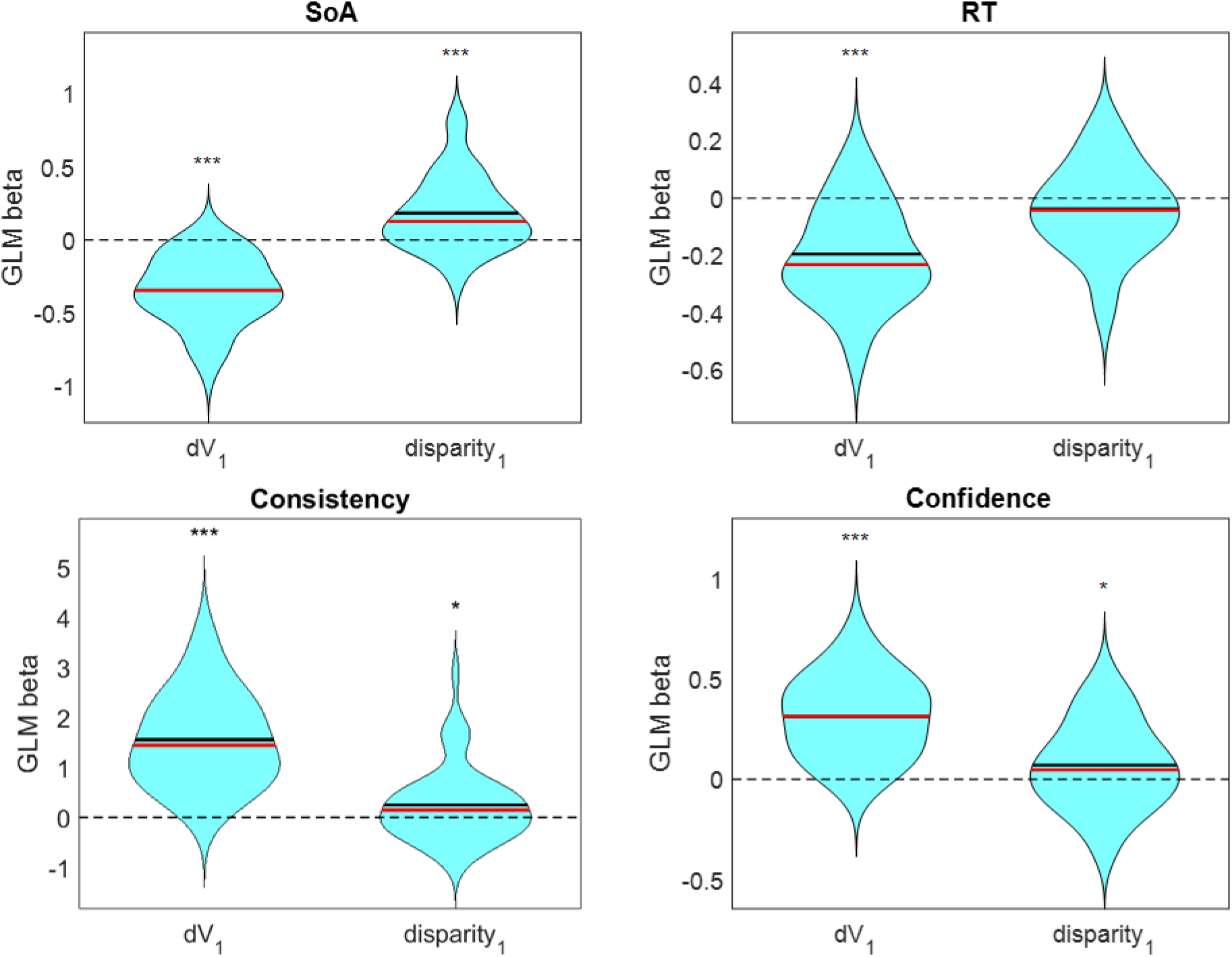
Impact of dV_1_ and disparity_1_ on SoA (top left), RT (top right), consistency (bottom left), and confidence (bottom right). Violin plots represent cross-participant distributions of GLM beta weights; black lines represent cross-participant mean values, red lines represent cross-participant median values; *** *p* < .001, ** *p* < .01, * *p* < .05.

#### Coherence Shifts in Attribute Values

The next set of analyses focused on coherence shifts at the level of the individual attributes, pleasure and nutrition. SoA values at the attribute level (SoA_P_ and SoA_N_ for pleasure and nutrition, respectively) were calculated in the analogous manner as overall SoA: in terms of the magnitude of the shift between the initial and final ratings of attribute values in the direction favoring the winner of the choice. Similarly, we defined the value difference for each attribute separately. For the initial ratings, these differences are termed dP_1_ and dN_1_, respectively.

We first checked to see if there was a reliable positive average SoA_P_ and SoA_N_ across trials. We observed a reliable SoA_P_ across all items (cross-participant mean of within-participant median SoA_P_ = 1.04, *p* = .025). We observed a smaller and less reliable SoA_N_ across all items (cross-participant mean of within-participant median SoA_N_ = 0.56, *p* = .108). (See the Supplementary Material for more details on SoA_P_ and SOA_N_.)

We then tested our predictions that dP_1_ would predict SoA_P_ and that dN_1_ would predict SoA_N_ using GLM regression. Across participants, both beta weights were negative and significant (for dP_1_ as a predictor of SoA_P_, mean beta = −0.210, *p* < .001; for dN_1_ as a predictor of SoA_N_, mean beta = −0.124, *p* < .001) (see Figure 5). The larger beta value for pleasure than nutrition likely reflects the greater subjective importance of the former attribute. Recall that dP and dN were coded as the attribute ratings for the higher-valued option minus the attribute ratings for the lower-valued option (based on pre-choice overall value ratings). These results imply that the attribute ratings shifted in favor of the chosen option, and to a larger extent if the chosen option initially rated poorly on that attribute (relative to the rejected option). If this attribute-specific SoA was merely a statistical artifact, we would expect the same regression results when using the absolute value of dP or dN (because the distinction of chosen versus rejected would no longer be relevant). We repeated the same regressions using |dP_1_| and |dN_1_| instead of dP_1_ and dN_1_. Across participants, the beta weight for |dP_1_| was negative and significant (mean beta = −0.234, *p* < .001), but the beta weight for |dN_1_| was not significant (mean beta = −0.031, *p* = .169; see Figure 5). The fact that the beta weights for dP_1_ and for |dP_1_| were similar likely reflects the fact that the pleasure rating for the chosen option was usually higher than for the rejected option (cross-participant median portion of trials = 0.72), whereas this was less frequently true for the nutrition rating (cross-participant median portion of trials = 0.50). We also assessed whether dP_1_ and dN_1_ would predict overall SoA. Across participants, neither beta weight was significant (dP_1_ *p* = .156, dN_1_ *p* = .105; see Figure 6). This suggests that overall SoA was not simply an affine combination of SoA_P_ and SoA_N_, and that coherence shifts occurred independently within the specific attribute dimensions.

**Figure 5.**
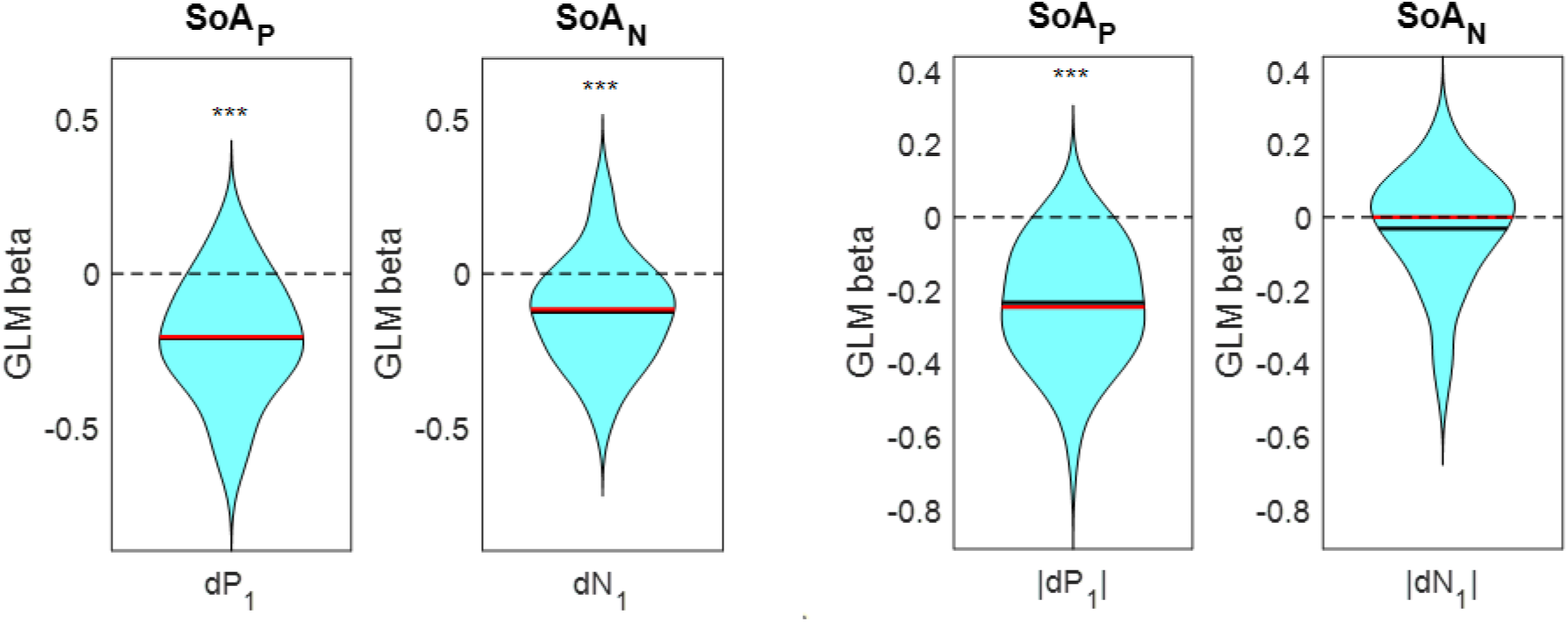
Impact of dP_1_ on SoA_P_ and dN_1_ on SoA_N_ (left) and of |dP_1_| on SoA_P_ and |dN_1_| on SoA_N_ (right). The left panel demonstrates the impact of the relative attribute rating of the chosen option versus the rejected option; the right panel demonstrates the impact of attribute rating similarity, regardless of which item was chosen.

**Figure 6.**
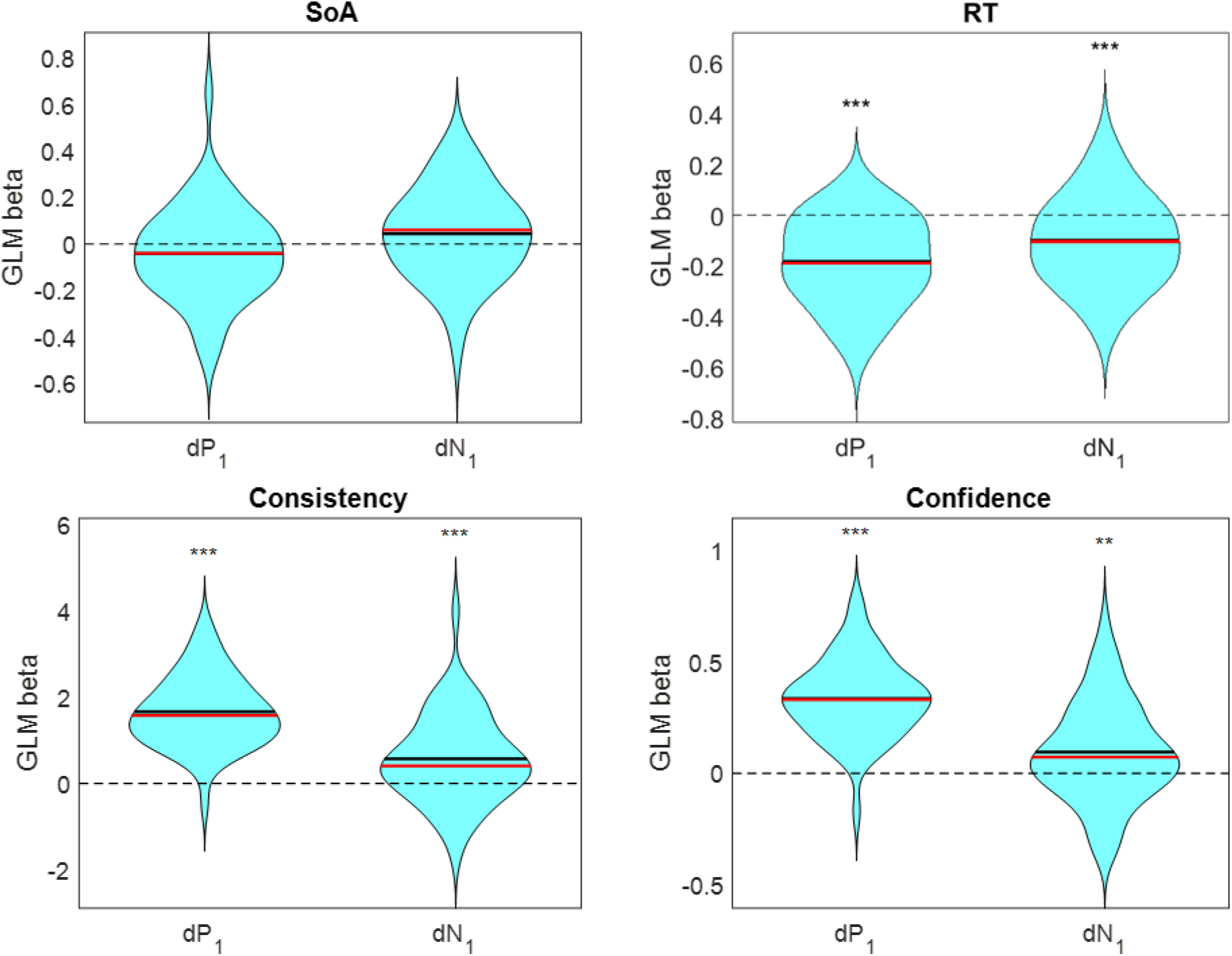
Impact of both dP_1_ and dN_1_ on SoA (middle left), on RT (middle right), on consistency (bottom left), and on confidence (bottom right). Violin plots represent cross-participant distributions of GLM beta weights; black lines represent cross-participant mean values, red lines represent cross-participant median values; *** *p* < .001, ** *p* < .01, * *p* < .05.

We then ran GLM regressions of both dP_1_ and dN_1_ on choice consistency, on choice confidence, and on RT. Across participants, beta weights for consistency were positive and significant for both independent variables (mean dP_1_ beta = 1.663, *p* < .001; mean dN_1_ beta = 0.566, *p* < .001), beta weights for confidence were positive and significant for both independent variables (mean dP_1_ beta = 0.338, *p* < .001; mean dN_1_ beta = 0.096, *p* = .004), and beta weights for RT were negative and significant for both independent variables (mean dP_1_ beta = −0.182, *p* < .001; mean dN_1_ beta = −0.100, *p* < .001; see Figure 6). When post-choice instead of pre-choice attribute ratings were used as predictors of consistency, the beta weight for dP_2_ was reliably greater than that for dP_1_ (*p* = .003, one-sided *t*-test), but that for dN_2_ did not differ from that for dN_1_ (*p* = .554, one-sided *t*-test). When post-choice attribute ratings were used as predictors of confidence, the beta weight for dP_2_ did not differ from that for dP_1_ (*p* = .754, one-sided *t*-test), nor did the beta weight for dN_2_ differ from that for dN_1_ (*p* = .934, one-sided *t*-test). When post-choice ratings were used as predictors of RT, the beta magnitude for dP_2_ was reliably greater than that for dP_1_ (*p* = .072, one-sided *t*-test), but that for dN_2_ did not differ from that for dN_1_ (*p* = .961, one-sided *t*-test).

## Experiments 2-5

Experiments 2-5 were designed to test the hypothesis that tasks other than explicit choice are also able to trigger coherence shifts. The general materials and design were basically the same as those of Experiment 1, except that ratings were obtained in three phases: initial ratings, ratings after a non-choice task, and finally ratings after a choice. As in Experiment 1, the final choice defined the winning option with respect to which all SoA measures were defined.

### Method

#### Participants

A total of 63 people participated in Experiment 2 (31 female; age: mean = 41 years, SD = 8, range 26-50). A total of 68 people participated in Experiment 3 (45 female; age: mean = 42 years, SD = 8, range 19-50). A total of 67 people participated in Experiment 4 (37 female; age: mean = 39 years, SD = 8, range 25-50). A total of 69 people participated in Experiment 5 (41 female; age: mean = 41 years, SD = 7, range 27-50). All participants were MTurk “masters”, residents of the United States or Canada, and native English speakers. Each participant received a payment of either $7.50 or $9 (increased for Experiments 4-5) as compensation for approximately one hour of time.

As in Experiment 1, we filtered our data for participants who might have performed the tasks carelessly. For each experiment, we calculated the Spearman correlation between first and last ratings, within participant, and also used GLM logistic regression of value ratings on choice to calculate the slope coefficient for each participant (separately using first and last ratings). Using these two measures, we deemed anyone with scores outside a cutoff (median + 3 * median average deviation) to be an outlier, and removed them from our analyses. After removing outliers, the number of participants whose data was used in analyses was 44 for Experiment 2, 48 for Experiment 3, 51 for Experiment 4, and 55 for Experiment 5.

#### Materials

For Experiments 2-5, we selected 60 stimuli (identical for all participants) from the full set of snacks (each presented as a digital image). The experimental set consisted of the 30 choice pairs (identical for all participants) that generated the highest overall SoA across participants in Experiment 1.

#### Design and Procedure

The basic design was very similar to that of Experiment 1, but with three phases of ratings (rather than two). After pre-exposure to all individual snacks, participants gave their initial ratings (phase 1) of overall value and of attribute values. This was followed by a non-choice task related to the 60 items, after which another set of ratings was obtained (phase 2). The choice task was then administered in the same way as in Experiment 1, and a final set of ratings was obtained (phase 3). Prior to administering the choice task, participants were not informed that any task would require a choice between options.

The pre-exposure, rating, and choice tasks were identical in format to those in Experiment 1. Across Experiments 2-5, the non-choice task varied from more to less “choice-like”. In Experiments 2-4, the 30 pairs of stimuli were displayed on the screen, one pair at a time, in a sequence randomized across participants (hence the two items were visible for potential comparison). These item pairs were the same as those that would be used in the choice task, except for the presentation sequence. Each individual item occurred in a single comparison pair. At the onset of each trial, a fixation cross appeared at the center of the screen for 750 ms. Next, a pair of images of food items appeared on the screen, one left and one right of center. In Experiment 2, participants responded to the question, “How similarly would you like these as daily snacks?” using a dropdown menu. The response choices were, “I like them {totally equal, very similar, fairly similar, slightly similar, totally different} amounts.” In Experiment 3, participants responded to the question, “How similar are these?” using a dropdown menu. The response choices were {totally similar, very similar, fairly similar, slightly similar, totally different}. Thus, whereas the similarity judgment made in Experiment 2 referred explicitly to “liking” of snacks, the similarity judgment in Experiment 3 made no reference to any sort of value. In Experiment 4, a green arrow appeared above one of the images on each trial (randomized between left and right), indicating which item the participants were to assess. Participants responded to the question, “When would you prefer to eat this?” using a dropdown menu. The response choices were {morning, afternoon, evening}. This task did not require participants to compare the two items in a pair, but did not prevent them from making a comparison.

In Experiment 5, in contrast to Experiments 1-4, the options were presented one at a time (rather than as pairs), so that a direct comparison of two items was impossible. Participants responded to the identical question as in Experiment 4 (i.e., time-of-day preference for a single snack). In all experiments, participants could revise their response as many times as they liked before finalizing it by clicking the “enter” button and proceeding to the next screen. No time limits were imposed for any of the constituent tasks or for the overall experiment.

### Results and Discussion

Experiments 2-5 included three phases of ratings. We will refer to measures of value difference (overall and for individual attributes) calculated using the initial round of ratings (prior to the non-choice task) with the subscript 1 (e.g., dV_1_), those based on the intermediate round of ratings (following the non-choice task but prior to the choice task) with the subscript 2, and those calculated using the final round of ratings (following the choice task) with the subscript 3. For SoA, subscript 1 is used for those measures calculated from pre- to post-non-choice (i.e., from phase 1 to phase 2). The subscript 2 is used for those measures calculated from post-non-choice to post-choice (i.e., from phase 2 to phase 3). For all measures of SoA, the winning choice is defined by the option eventually selected during the choice task. Critically, SoA_1_ values reflect coherence shifts that occurred *before* the choice task was even administered. Positive values of SoA_1_, SoAP_1_, and SoAN_1_ will thus indicate changes in the direction of the eventual winner triggered by a task that did not itself require making a choice. Similarly, any impact of non-choice tasks on eventual choice confidence precedes the posing of a choice task.

#### Coherence Shifts in Overall Value During Non-Choice Tasks

Our main goal in Experiments 2-5 was to assess whether each of the effects related to choice that were observed in Experiment 1 would also be triggered by our novel non-choice task design. Indeed, we found a reliable positive level of SoA_1_ when we examined the change in overall value ratings before and after the non-choice tasks in each of Experiments 2-5. As anticipated, the magnitude of SoA_1_ in these experiments was largest in Experiment 2 (mean of median SoA_1_ = 3.19, *p* < .001), followed by Experiment 3 (mean of median SoA_1_ = 2.22, *p* = .019), followed by Experiment 4 (mean of median SoA_1_ = 1.73, *p* = .008), with Experiment 5 being the lowest (mean of median SoA_1_ = 1.18, *p* = .068). These results are summarized in Table 1 below. Note the clear decreasing pattern of SoA_1_ as the non-choice task moves from more to less “choice-like”.

**Table 1:** Across the different experiments, the cross-participant mean of within-participant SoA was positive and significant, and exhibited a downward-trending pattern as the non-choice task became less comparative.

|  | SoA<sub>1</sub> |
| --- | --- |
| Experiment 2 (similarity - value) | 3.193 |
| Experiment 3 (similarity - general) | 2.219 |
| Experiment 4 (time of day - pairs) | 1.726 |
| Experiment 5 (time of day - singles) | 1.182 |

The cross-experiment gradient of SoA_1_ supports our claim that a major cause of SoA is cognitive contemplation of the options, rather than being either a mere statistical artifact or triggered only by an explicit choice (as cognitive dissonance theorists would hold). To provide additional support for our claim, we ran GLM regressions of dV_1_, dV_2_, and dV_3_ (separately) on eventual choice consistency, on choice confidence, and on RT. All beta weights were significant and in the predicted direction (see Table 2; *p* < .001 for all beta weights). More interestingly, in each experiment the magnitude of the beta weights for dV on each of the dependent variables increased from phase 1 to 2 and from phase 2 to 3. Not all of the differences in beta weights were statistically significant (see Figure 7), but the trend was systematic and robust across dependent variables and across experiments (see Table 2 and Figure 7). Thus, value differences after coherence shifts (both between phases 1 and 2, and between phases 2 and 3) were better predictors of choice consistency, choice confidence, and RT than were the initial value differences. Notably, this pattern would not be expected if the coherence shifts were due to post-choice cognitive dissonance resolution or were solely a statistical artifact of repeated ratings.

**Figure 7.**
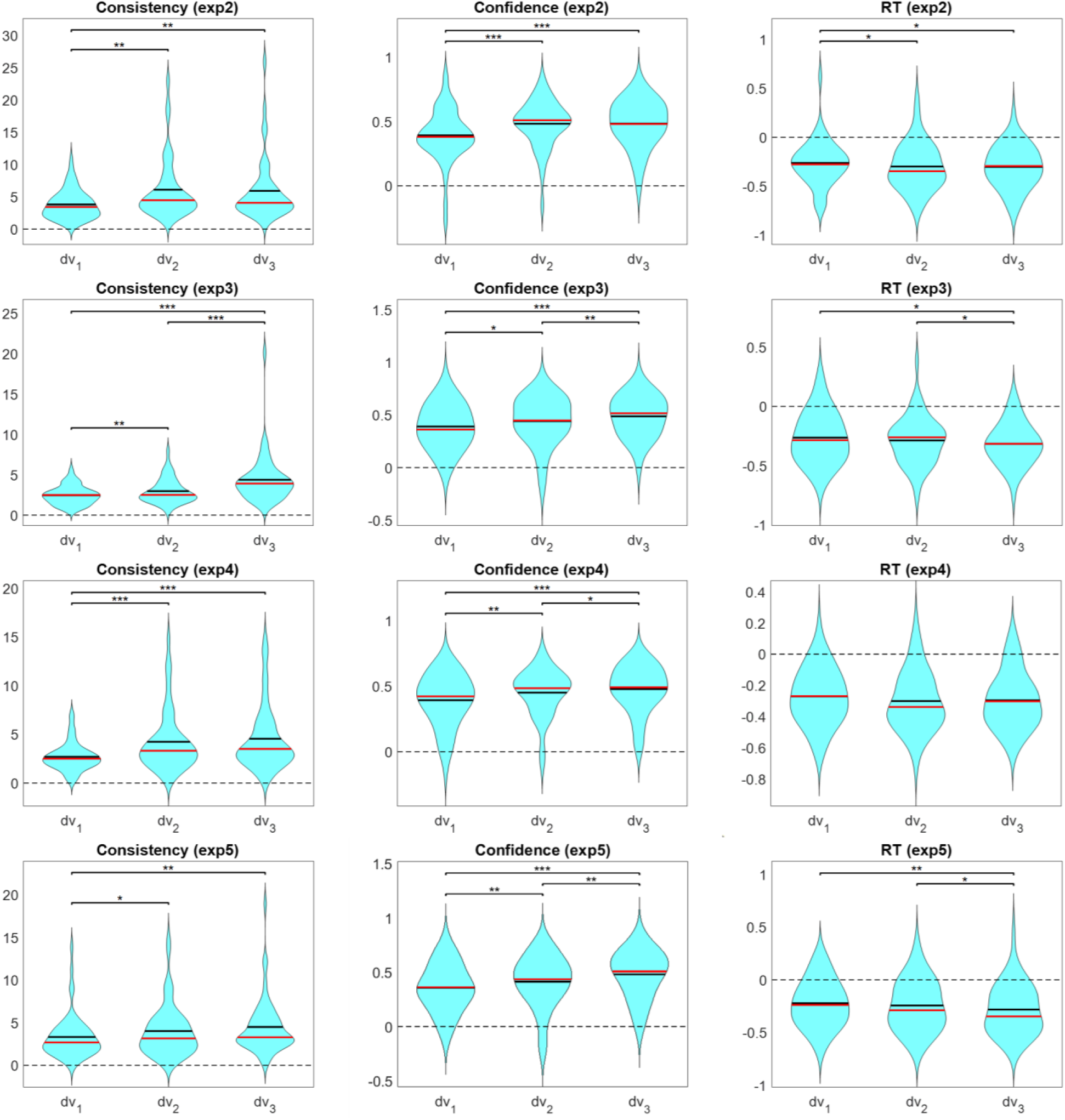
Impact of dV on consistency (left panels), on confidence (middle panels), and on RT (right panels), separately for each round of ratings. Each row of plots is for a different experiment. Violin plots represent cross-participant distributions of GLM beta weights; black lines represent cross-participant mean values, red lines represent cross-participant median values; *** *p* < .001, ** *p* < .01, * *p* < .05.

**Table 2:** For Experiments 2-5, the impact of dV on consistency, on confidence, and on RT increased from first to second to third rating rounds.

| **CHOICE CONSISTENCY** | $\beta_{dv1\_Ch}$ | $\beta_{dv2\_Ch}$ | $\beta_{dv3\_Ch}$ |
| --- | --- | --- | --- |
| Experiment 2 (similarity - value) | 3.810 | 6.107 | 5.924 |
| Experiment 3 (similarity - general) | 2.462 | 2.971 | 4.385 |
| Experiment 4 (time of day - pairs) | 2.698 | 4.231 | 4.549 |
| Experiment 5 (time of day - singles) | 3.324 | 4.010 | 4.492 |
| **CHOICE CONFIDENCE** | $\beta_{dv1\_CC}$ | $\beta_{dv2\_CC}$ | $\beta_{dv3\_CC}$ |
| Experiment 2 (similarity - value) | 0.393 | 0.484 | 0.485 |
| Experiment 3 (similarity - general) | 0.391 | 0.443 | 0.488 |
| Experiment 4 (time of day - pairs) | 0.393 | 0.451 | 0.478 |
| Experiment 5 (time of day - singles) | 0.355 | 0.413 | 0.479 |
| **RESPONSE TIME** | $\beta_{dv1\_RT}$ | $\beta_{dv2\_RT}$ | $\beta_{dv3\_RT}$ |
| Experiment 2 (similarity - value) | -0.263 | -0.299 | -0.303 |
| Experiment 3 (similarity - general) | -0.264 | -0.287 | -0.316 |
| Experiment 4 (time of day - pairs) | -0.271 | -0.301 | -0.297 |
| Experiment 5 (time of day - singles) | -0.222 | -0.244 | -0.282 |

The results presented above suggest that ratings provided later in the experiment are more in line with the “true” evaluations revealed by the explicit choice task. This increase in precision should be closely linked to SoA, since SoA_1_ and SoA_2_ are basically dV_2_-dV_1_ and dV_3_-dV_2_, respectively (although they are not mathematically identical, because preference reversals can increase the magnitude of SoA). To demonstrate conclusively the relationship between SoA and the dependent variables of interest, we performed GLM regressions of dV1, SoA1, and SoA2 on choice confidence and on RT.^1^ Across all experiments, all beta weights were significant and in the expected direction (positive for confidence, negative for RT; see Figure 8).

**Figure 8.**
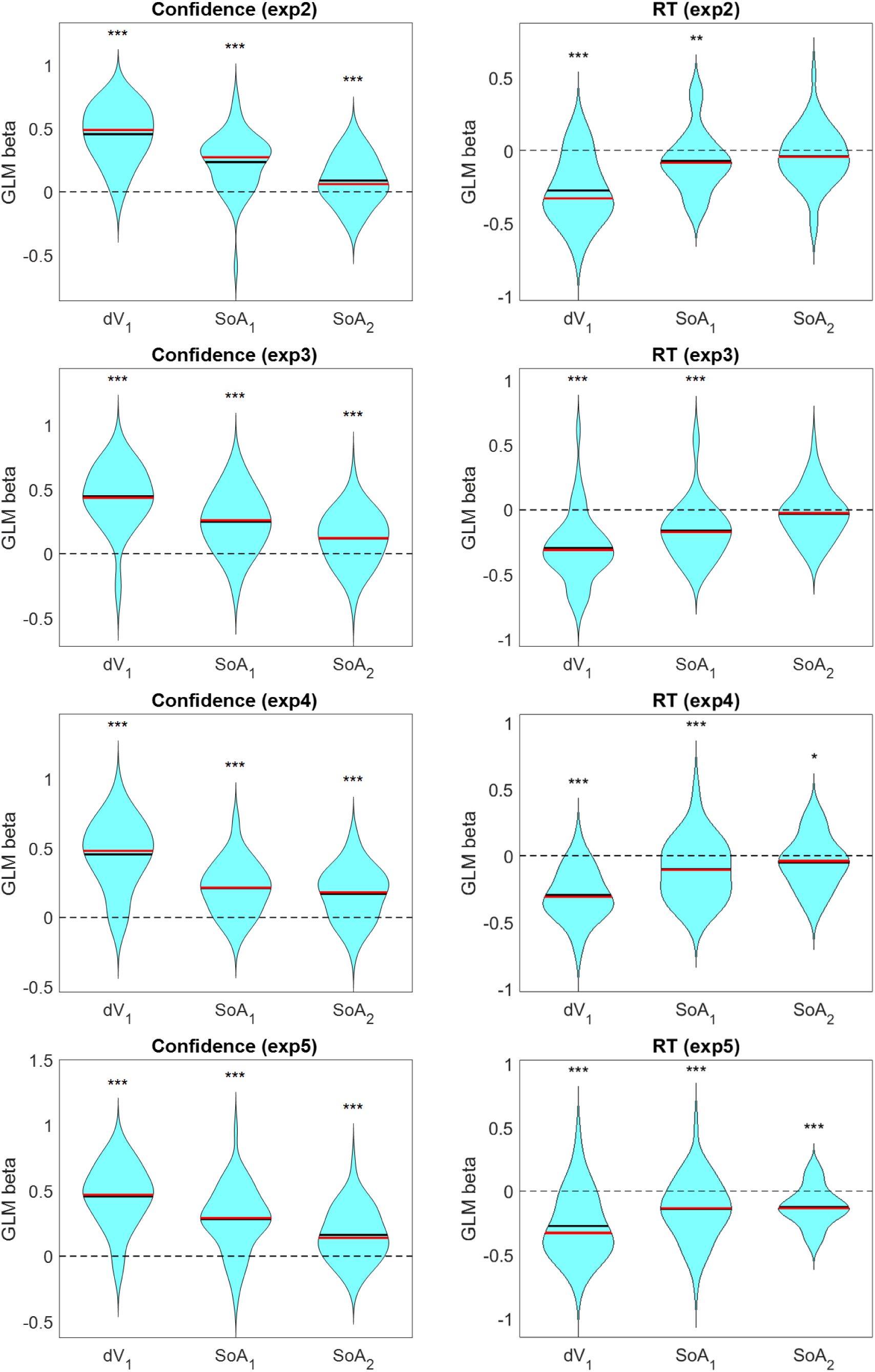
Impact of dV, SoA_1_, and SoA_2_ on confidence (left panels) and on RT (right panels). Each row of plots is for a different experiment. Violin plots represent cross-participant distributions of GLM beta weights; black lines represent cross-participant mean values, red lines represent cross-participant median values; *** p < .001, ** p < .01, * p < .05.

Finally, we verified that our additional regression analyses from Experiment 1 replicated across each of the other experiments. Specifically, we ran GLM regressions of dV_1_ and SoA_1_ on RT and on confidence. We also ran GLM regressions of dV_1_ and disparity_1_ on SoA_1_, on RT, and on confidence. Across participants, all beta weights were significant and in the predicted direction (see Supplementary Material, section Statistical Summary).

### General Discussion

The present study provides a number of important findings concerning the role of coherence shifts in the construction of preferences. When required to choose between two options (snacks), each differing in two attributes that determine value (pleasure and nutrition), people’s assessments of value shifted from pre- to post-choice in the direction that spread the alternatives further apart so as to favor the winner, thereby increasing confidence in the choice. This shift was observed not only for ratings of overall value, but also for each of the two individual attributes that determined value. Moreover, the magnitude of the coherence shift increased with the difficulty of the choice as measured by the difference in initial ratings of overall value for the two options. Coherence shifts were in turn predictive of increased choice confidence and decreased RT, with each of these dependent variables being more accurately predicted by value difference measured by ratings obtained after the choice.

We also found that coherence shifts are predicted not only by the overall value difference between options, but also by the pattern of attribute composition across options (which we believe is a novel contribution to the literature). We introduced a formal measure of attribute *disparity* (Equation 1), reflecting the degree to which individual attributes “disagree” with one another as to which option is superior. After accounting for the impact of overall value difference, the magnitude of the coherence shift was also positively correlated with disparity. Moreover, coherence shifts associated with disparity increased confidence and decreased RT in the eventual choice.

Experiments 2-5 provided evidence that coherence shifts are driven by refinements of the mental representations of the options, and not merely by post-choice adjustments (e.g., cognitive dissonance reduction) or statistical noise (i.e., a regression to the mean effect across repeated evaluations). Specifically, Experiment 2 demonstrated that an active comparison of option values is likely the core component of the decision process that drives the coherence shifts typically observed in a choice task. To isolate the impact of the comparison process, Experiment 2 used a three-phase design in which value ratings were obtained at the outset, after a value comparison task, and finally after an actual choice task. Critically, the comparison task was administered prior to informing the participants that any choice would be required. Instead, they were simply asked to rate the similarity of values for two snacks. This comparison task generated the same qualitative pattern of changes in value ratings and in consistency, confidence, and RT in the eventual choice, even though the similarity judgments entirely preceded the actual choice task. These findings support the hypothesis that active comparison of values is a core component of the decision process during which perceived values are shifted so as to more sharply distinguish the items being compared.

Moreover, Experiments 3-5 demonstrated that an active comparison of option values is not even necessary to cause coherence shifts. Specifically, Experiment 3 replaced similarity judgments based on value comparison (Experiment 2) with a more generic similarity judgment not explicitly linked to value. While it is possible that some participants compared the snacks in terms of value, it is likely that non-value aspects (e.g., size, shape, and color) were also considered. The generic similarity task nevertheless triggered coherence shifts in the eventual choices (between the same pairs of items for which similarity judgments had been made). This finding suggests that any additional processing directed at the options on offer, even if not explicitly cued towards value, leads to a refinement of the mental representations of the options. This refinement automatically brings the benefit of more accurate valuations when solicited in subsequent rating tasks.

In Experiment 4, we replaced the comparison task with an even more generic task. Here, we presented the same pairs of snacks as in the eventual choice task, but asked participants to ignore one option while deciding at what time of day they would prefer to eat the cued option. Notably, this task requires absolutely no comparison between the options. Nevertheless, it appears to have caused some degree of coherence shifts (though less than in the explicit comparison tasks). This finding suggests that when presented with pairs of categorically similar options (in this case, snacks), the brain may automatically perform some initial rough comparison of their values, even when value is not task-relevant and the overt task concerns just one of the paired items. This interpretation is consistent with previous work showing that the brain automatically encodes the relative value of options even when the task is unrelated to value (Grueschow et al., 2015).

In Experiment 5, we used the same time-of-day judgment as in Experiment 4 but with displays showing a single item at a time, thus precluding even the possibility of a comparison between options while performing the task. We still observed some positive level of coherence shifts in Experiment 5 (though with a lower magnitude than in any of the other experiments). This finding suggests that consideration of individual snacks in isolation (rather than in contrast to other snacks) can also lead to refinement of their mental representation. This result is consistent with previous work showing that the brain automatically encodes value of individual options even when the task is unrelated to value (Lebreton et al., 2009). The automatic value signal for a particular option may adjust the latent value estimate in the mind of the decision maker, altering the probability that that option will later be chosen in a subsequent choice task.

Taken together, our results cast serious doubt on several alternative explanations of observed coherence shifts. Accounts based on post-choice resolution of cognitive dissonance are unable to explain why we observed coherence shifts in a variety of non-choice situations. Furthermore, dissonance reduction cannot explain the observed gradient in coherence shifts across the different types of task. Our data also rule out suggestions that coherence shifts are solely due to statistical artifacts. Under such an account, the value representations of options will not change as a function of intervening tasks. Accordingly, any observed relationship between value difference (dV) and choice behavior (consistency, confidence, RT) should be equally well predicted using ratings from any of the experimental phases. The fact that we observed a clear increase in the explanatory power of dV on all dependent variables from the first to second to third phase makes it highly unlikely that the evolution of dV across phases (via SoA_1_ and SoA_2_) was mere statistical noise.

Lee and Daunizeau (2020b) have recently proposed a computational account of why and how coherence shifts can be generated during intra-decisional processing. According to their metacognitive control of decision-making (MCD) model, decision makers should exert mental effort only to the degree necessary to reach some subjective threshold of confidence that the chosen option is indeed the best of the candidate set. People process additional information about choice options up until the point at which the options are sufficiently distinguishable to provide a satisfactory level of confidence in the emerging choice. The MCD model can account for empirical evidence that people change their subjective value estimates when required to choose between options that they initially estimated to have similar values, and that this change in value correlates with both choice confidence and response time. Notably, while the authors presented this model specifically in the context of active deliberation during an explicit choice, it could likely be extended to account for the impact of information processing in other tasks (as in Experiments 2-5). For example, contemplation during any task may lead decision makers closer to some sort of “ground truth” (e.g., if they consider more attributes that they previously ignored), leading to ratings that are both more accurate and more precise (and thus the option with the “true” higher value would be more likely to be chosen in a subsequent choice task). Beyond that, a head-to-head comparison of options (e.g., during a choice task) may magnify the spreading effect, because the relative values of the attributes (between options) may be more salient than the values of each option evaluated in isolation.

The behavioral evidence concerning intra-decisional processing provided by the present study fits well with the picture emerging from studies of the emergence of decisions at the neural level. It has been shown that the brain computes attribute values separately and then integrates them before making a choice (Lim, O’Doherty, & Rangel, 2013). Other neural evidence indicates that the assessment of individual attributes, as well as their relative weights on the final choice, evolve during the decision process (Hunt et al., 2014). Löffler, Haggard, and Bode (2020; also Voigt et al., 2019) have provided compelling neuroimaging evidence that shifts in valuation that support the eventual choice occur prior to the choice response. Future studies using multiple methodologies will hopefully provide a deeper understanding of the timing of coherence shifts at the attribute level. In particular, it would be informative to examine how activity patterns in key brain regions vary across different sorts of choice or non-choice tasks, in which the refinement of value representations seems to take place to varying degrees.

It has been proposed that CIPC is mediated by the strength of memory encoding for individual options. Several studies (Botvinik-Nezer et al., 2019; Bakkour, et al., 2017; Schonberg et al., 2014) showed that enhanced processing of snack food options (based on response cueing for specific options during sequential passive viewing) led to better option recall for those options, and that (regardless of cueing) better recall was associated with a higher probability of being chosen (when paired with alternatives of similar subjective value). This suggests that the more precise value representations associated with cued (or otherwise better remembered) options enhanced the apparent value of those options. Previous work has also shown that CIPC only occurs for options that are remembered well (Chammat et al., 2017). Other previous work has demonstrated stronger memory encoding for options of higher value (Shohamy & Adcock, 2010; Miendlarzewska et al., 2015), which could play a causal role in the spreading of alternatives phenomenon (higher-valued options would be more likely to be chosen, and also more precisely encoded). Schonberg and Katz (2020) proposed a brain network potentially responsible for CIPC, based on selective attention. Further studies are needed to better understand the relationship between value, memory, and preference change.

Another useful direction for future research would be to collect self-reported estimates of attribute-specific importance weights, rather than relying on statistical regression to infer these weights. Recall that in the present experiments, final ratings were obtained using all individual items after completing all choices between paired options. Each individual item occurred in only one choice trial; hence any value shift for an item can be attributed to that one specific choice. However, the two basic attributes were likely assessed on all choice trials, and hence their relative importance would have been pushed in different directions on different trials (presumably canceling out any coherence shifts due to trial-by-trial changes in attribute importance weights). A different paradigm would be required to meaningfully assess coherence shifts in attribute importance. Such a paradigm could also be used to assess whether the magnitude of coherence shifts along a particular attribute dimension is systematically correlated with the self-reported importance of that attribute.

## Acknowledgements

Preparation of this paper was supported by NSF Grant BCS-1827374. We thank Dan Simon for comments on an earlier draft of this paper.

## Supplementary Material

### Data Quality Checks

Before conducting our main analyses, we performed a number of basic quality checks to verify that our data were reliable. First, we assessed the test-retest reliability of overall value ratings. For each participant, we measured the correlation between first (R1) and second (R2) ratings (both experiments) and between first and third (R3) ratings (Experiments 2-5), across items. We found that ratings were generally consistent (in Experiment 1, median Spearman’s rho for R1-R2 = 0.830; in Experiment 2, for R1-R2 = 0.857, and for R1-R3 = 0.840). We then performed the same tests for pleasure ratings, and found that ratings were generally consistent (in Experiment 1, median Spearman’s rho for R1-R2 = 0.835; in Experiment 2, for R1-R2 = 0.877, for R1-R3 = 0.865). We then performed the same tests for nutrition ratings, and found that ratings were generally consistent (in Experiment 1, median Spearman’s rho for 1 R1-R2 = 0.885; in Experiment 2, for R1-R2 = 0.886, for R1-R3 = 0.860). These correlations are presented in Figure S1. The data for Experiments 3-5 were similar.

Beyond test-retest reliability, we also showed that the distributions for overall ratings, for pleasure ratings, and for nutrition ratings did not change between experimental task sections (see Figure S2). This is important, because if participants changed how they used the rating scale as they rated more options, a mere effect of habituation to the rating scale could have caused the observed spreading of alternatives effect. However, we found no evidence that this was the case in any of the experiments.

Next, we checked whether the range of values for both attributes (pleasure and nutrition) was the same for each participant. This would be important because we relied on GLM regression to estimate the attribute importance weights upon which our disparity calculations were based. If the range of one attribute was substantially different than that for the other, then the beta weights obtained by GLM regression might not actually be indicative of how much each attribute contributes to overall value. We found that most participants used the full range of possible values for both pleasure and nutrition, and that the full pleasure-nutrition space was sufficiently covered (see Figure S3). Our techniques reported in the main manuscript were thus acceptable.

Because many of our analyses relied on the dV and disparity variables, we checked to ensure that each of these variables was sufficiently well distributed across a wide-enough range of values. We found that the distributions were similar, and that the dV-disparity space was sufficiently covered (see Figure S4).

### Attribute Importance Weights

Because our experimental design has never before been used (to our knowledge) with attribute-specific ratings, we wanted to verify that the attribute ratings we collected were truly related to overall value as we expected them to be. To do this, we first regressed (GLM) the pleasure and nutrition attribute ratings (R1) of the items onto the overall value ratings, separately for each participant. Across participants in each experiment, there was a reliable contribution of both attributes to value ratings (Experiment 1: mean pleasure beta = 0.653, *p* < .001; mean nutrition beta = 0.157, *p* < .001; Experiment 2: mean pleasure beta = 0.660, *p* < .001; mean nutrition beta = 0.118, *p* < .001). These effects were stronger when we used the second round of attribute and value ratings (R2) instead of the first (Experiment 1: mean pleasure beta = 0.763, *p* < .001; mean nutrition beta = 0.192, *p* < .001; Experiment 2: mean pleasure beta = 0.733, *p* < .001; mean nutrition beta = 0.154, *p* < .001). The difference in the beta weights using R2 vs R1 ratings was significant for pleasure (Experiment. 1: *p* < .001; Experiment 2: *p* < .001), but only moderately significant for nutrition (Experiment 1: *p* = .096; Experiment 2: *p* = .029). For Experiment 2, these effects were stronger when we used the third round of attribute and value ratings (R3) instead of the second (mean pleasure beta = 0.780, *p* < .001; mean nutrition beta = 0.189, *p* < .001). The difference in the beta weights using R3 vs R1 ratings was significant for both pleasure (*p* = .039) and nutrition (*p* = .047). These results imply that participants had a more precise sense of their subjective valuations (in terms of attribute contribution to overall value) for the set of items after deliberation (for both similarity judgment and explicit choice) relative to before (see Figure S5). The data for Experiments 3-5 were similar to the data for Experiment 2.

### Spreading of Alternatives

In the Results section of the main manuscript, we report group averages for the mean and median within-participant spreading of alternatives (SoA). SoA was highly variable across trials for every participant, implying that much of the observed SoA effect in our data is driven by either noise or some unknown factor. Nevertheless, SoA is reliably greater than zero on average (both mean and median) for most participants. This holds true for both overall value SoA, and for attribute-specific SoA (SoA_P_ and SoA_N_; see Figure S6).

### Similarity Judgments

We realize that the phrasing of our similarity judgment question in Experiment 2 (“How similarly would you like these items as a daily snack?”) was a bit unnatural. It was therefore possible that participants might have been confused about how to properly respond to such a question. It was our intention that participants would assess the similarity of the options in terms of their subjective overall values. We thus checked to see how well the reported similarity judgments for the choice pairs correlated with their differences in overall value (as reported in R1). We found that, as expected, the two measures were highly negatively correlated across participants (mean Spearman’s rho = −0.410, *p* < .001; see Figure S7). Also as expected, the magnitude of the correlation increased when using post-judgment ratings (R2) (mean Spearman’s rho = −0.489, *p* < .001), and the difference in the two correlations was significant (*p* < .001).

### Statistical Summary

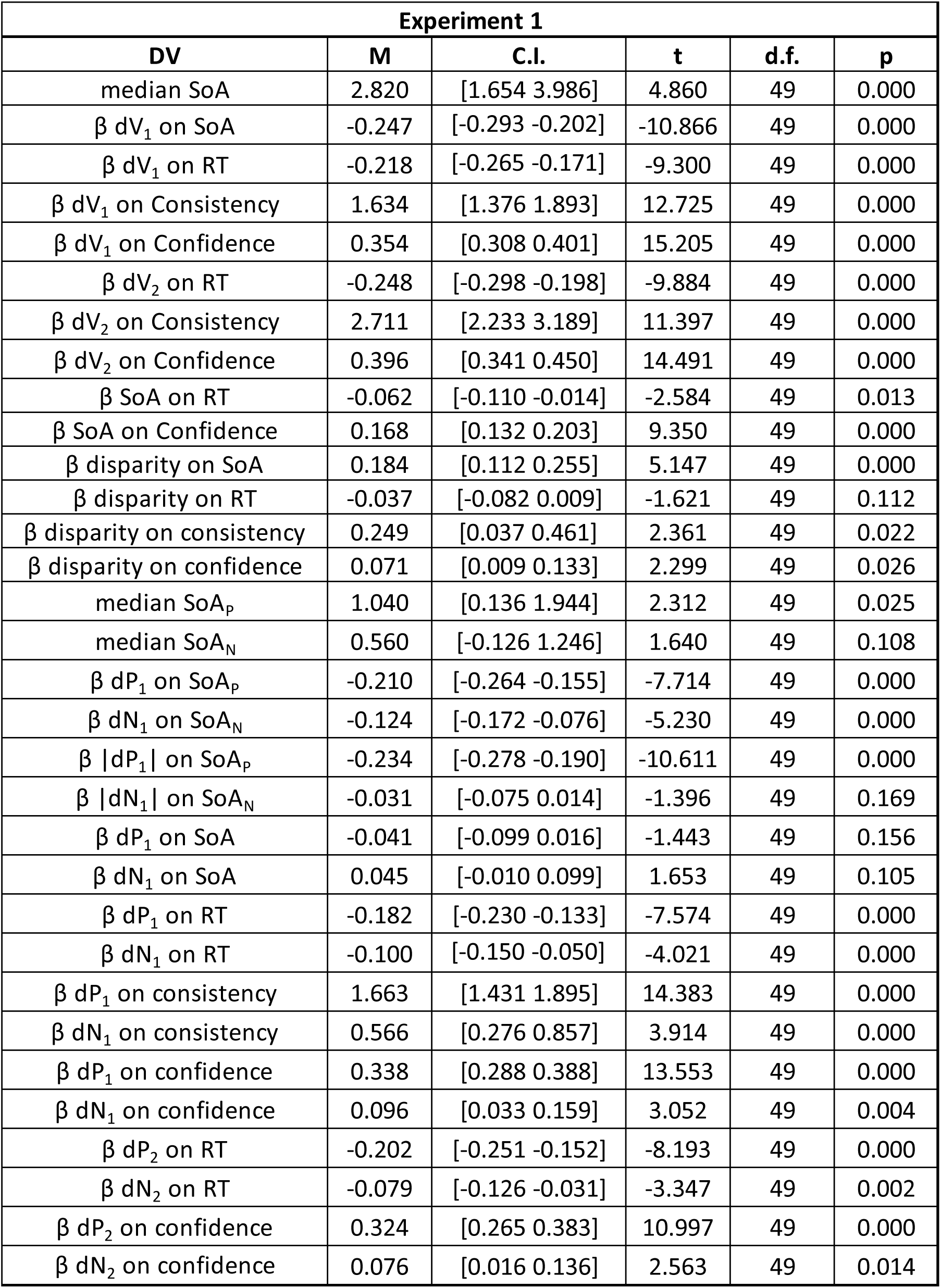

| Experiment 2 |  |  |  |  |  |
| --- | --- | --- | --- | --- | --- |
| DV | M | C.I. | t | d.f. | p |
| median SoA<sub>1</sub> | 3.193 | [1.955 4.431] | 5.201 | 43 | 0.000 |
| $\beta$ dV<sub>1</sub> on RT | -0.263 | [-0.334 -0.192] | -7.458 | 43 | 0.000 |
| $\beta$ dV<sub>1</sub> on Consistency | 3.810 | [3.154 4.466] | 11.712 | 43 | 0.000 |
| $\beta$ dV<sub>1</sub> on Confidence | 0.393 | [0.329 0.456] | 12.481 | 43 | 0.000 |
| $\beta$ dV<sub>2</sub> on RT | -0.299 | [-0.371 -0.227] | -8.325 | 43 | 0.000 |
| $\beta$ dV<sub>2</sub> on Consistency | 6.107 | [4.677 7.537] | 8.614 | 43 | 0.000 |
| $\beta$ dV<sub>2</sub> on Confidence | 0.484 | [0.428 0.539] | 17.439 | 43 | 0.000 |
| $\beta$ dV<sub>3</sub> on RT | -0.303 | [-0.367 -0.239] | -9.478 | 43 | 0.000 |
| $\beta$ dV<sub>3</sub> on Consistency | 5.924 | [4.453 7.394] | 8.124 | 43 | 0.000 |
| $\beta$ dV<sub>3</sub> on Confidence | 0.485 | [0.428 0.541] | 17.295 | 43 | 0.000 |
| $\beta$ SoA<sub>1</sub> on RT | -0.161 | [-0.229 -0.093] | -4.773 | 43 | 0.000 |
| $\beta$ SoA<sub>2</sub> on RT | -0.030 | [-0.089 0.029] | -1.017 | 43 | 0.315 |
| $\beta$ SoA<sub>1</sub> on Confidence | 0.250 | [0.181 0.320] | 7.286 | 43 | 0.000 |
| $\beta$ SoA<sub>2</sub> on Confidence | 0.119 | [0.058 0.180] | 3.957 | 43 | 0.000 |
| $\beta$ SoA on RT | -0.145 | [-0.206 -0.084] | -4.818 | 43 | 0.000 |
| $\beta$ SoA on Confidence | 0.210 | [0.144 0.276] | 6.385 | 43 | 0.000 |
| $\beta$ disparity on SoA | 0.350 | [0.270 0.429] | 8.896 | 43 | 0.000 |
| $\beta$ disparity on RT | -0.092 | [-0.152 -0.032] | -3.106 | 43 | 0.003 |
| $\beta$ disparity on confidence | 0.203 | [0.128 0.279] | 5.459 | 43 | 0.000 |

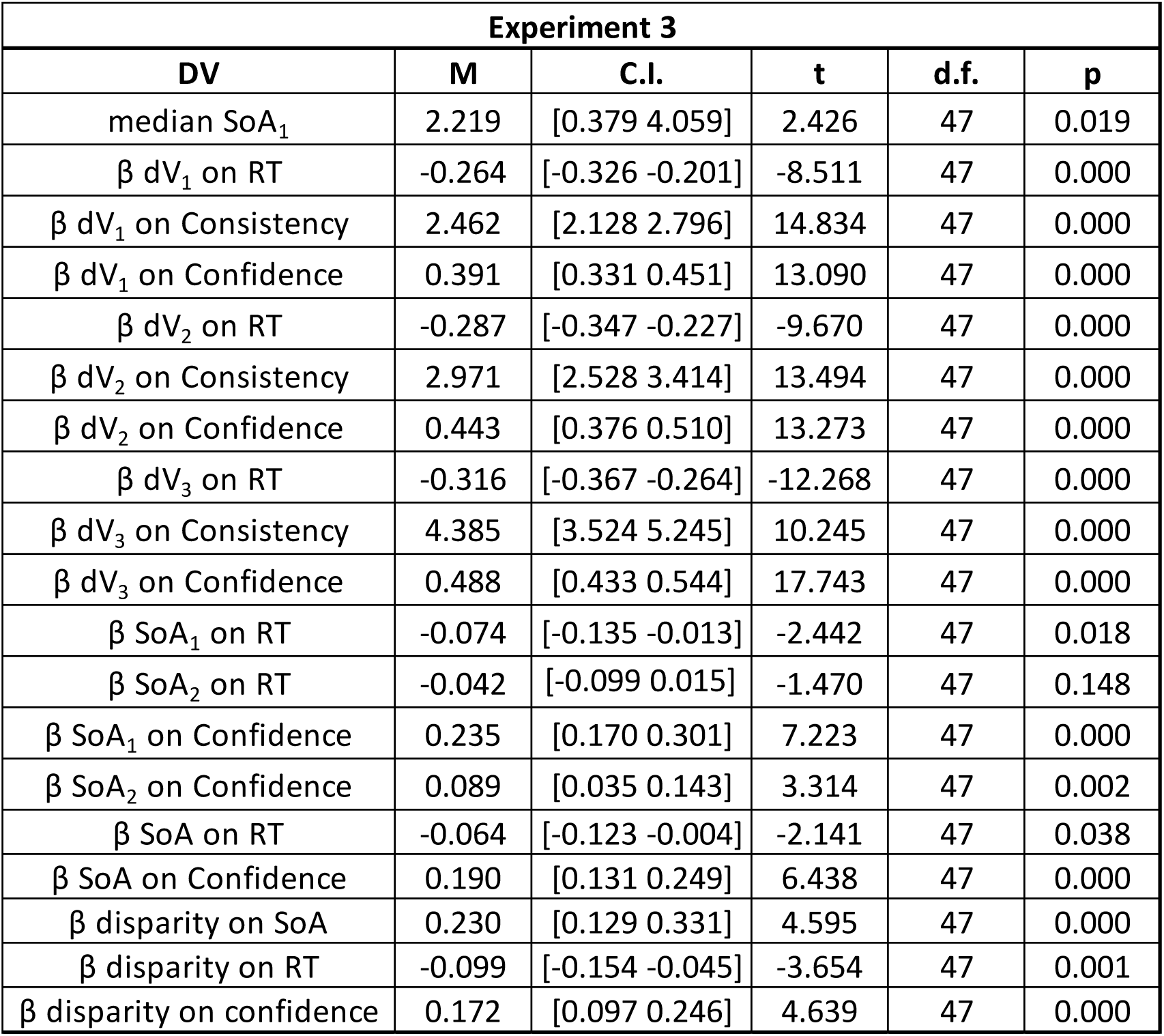

| Experiment 3 |  |  |  |  |  |
| --- | --- | --- | --- | --- | --- |
| DV | M | C.I. | t | d.f. | p |
| median SoA<sub>1</sub> | 2.219 | [0.379 4.059] | 2.426 | 47 | 0.019 |
| $\beta$ dV<sub>1</sub> on RT | -0.264 | [-0.326 -0.201] | -8.511 | 47 | 0.000 |
| $\beta$ dV<sub>1</sub> on Consistency | 2.462 | [2.128 2.796] | 14.834 | 47 | 0.000 |
| $\beta$ dV<sub>1</sub> on Confidence | 0.391 | [0.331 0.451] | 13.090 | 47 | 0.000 |
| $\beta$ dV<sub>2</sub> on RT | -0.287 | [-0.347 -0.227] | -9.670 | 47 | 0.000 |
| $\beta$ dV<sub>2</sub> on Consistency | 2.971 | [2.528 3.414] | 13.494 | 47 | 0.000 |
| $\beta$ dV<sub>2</sub> on Confidence | 0.443 | [0.376 0.510] | 13.273 | 47 | 0.000 |
| $\beta$ dV<sub>3</sub> on RT | -0.316 | [-0.367 -0.264] | -12.268 | 47 | 0.000 |
| $\beta$ dV<sub>3</sub> on Consistency | 4.385 | [3.524 5.245] | 10.245 | 47 | 0.000 |
| $\beta$ dV<sub>3</sub> on Confidence | 0.488 | [0.433 0.544] | 17.743 | 47 | 0.000 |
| $\beta$ SoA<sub>1</sub> on RT | -0.074 | [-0.135 -0.013] | -2.442 | 47 | 0.018 |
| $\beta$ SoA<sub>2</sub> on RT | -0.042 | [-0.099 0.015] | -1.470 | 47 | 0.148 |
| $\beta$ SoA<sub>1</sub> on Confidence | 0.235 | [0.170 0.301] | 7.223 | 47 | 0.000 |
| $\beta$ SoA<sub>2</sub> on Confidence | 0.089 | [0.035 0.143] | 3.314 | 47 | 0.002 |
| $\beta$ SoA on RT | -0.064 | [-0.123 -0.004] | -2.141 | 47 | 0.038 |
| $\beta$ SoA on Confidence | 0.190 | [0.131 0.249] | 6.438 | 47 | 0.000 |
| $\beta$ disparity on SoA | 0.230 | [0.129 0.331] | 4.595 | 47 | 0.000 |
| $\beta$ disparity on RT | -0.099 | [-0.154 -0.045] | -3.654 | 47 | 0.001 |
| $\beta$ disparity on confidence | 0.172 | [0.097 0.246] | 4.639 | 47 | 0.000 |

| Experiment 4 |  |  |  |  |  |
| --- | --- | --- | --- | --- | --- |
| DV | M | C.I. | t | d.f. | p |
| median SoA<sub>1</sub> | 1.726 | [0.475 2.976] | 2.771 | 50 | 0.008 |
| $\beta$ dV<sub>1</sub> on RT | -0.271 | [-0.322 -0.221] | -10.867 | 50 | 0.000 |
| $\beta$ dV<sub>1</sub> on Consistency | 2.698 | [2.294 3.102] | 13.400 | 50 | 0.000 |
| $\beta$ dV<sub>1</sub> on Confidence | 0.393 | [0.337 0.449] | 14.032 | 50 | 0.000 |
| $\beta$ dV<sub>2</sub> on RT | -0.301 | [-0.355 -0.247] | -11.285 | 50 | 0.000 |
| $\beta$ dV<sub>2</sub> on Consistency | 4.231 | [3.358 5.104] | 9.734 | 50 | 0.000 |
| $\beta$ dV<sub>2</sub> on Confidence | 0.451 | [0.400 0.501] | 17.831 | 50 | 0.000 |
| $\beta$ dV<sub>3</sub> on RT | -0.297 | [-0.348 -0.245] | -11.604 | 50 | 0.000 |
| $\beta$ dV<sub>3</sub> on Consistency | 4.549 | [3.618 5.479] | 9.818 | 50 | 0.000 |
| $\beta$ dV<sub>3</sub> on Confidence | 0.478 | [0.427 0.529] | 18.841 | 50 | 0.000 |
| $\beta$ SoA<sub>1</sub> on RT | -0.105 | [-0.168 -0.043] | -3.400 | 50 | 0.001 |
| $\beta$ SoA<sub>2</sub> on RT | -0.054 | [-0.110 0.002] | -1.951 | 50 | 0.057 |
| $\beta$ SoA<sub>1</sub> on Confidence | 0.212 | [0.159 0.265] | 8.045 | 50 | 0.000 |
| $\beta$ SoA<sub>2</sub> on Confidence | 0.172 | [0.119 0.224] | 6.569 | 50 | 0.000 |
| $\beta$ SoA on RT | -0.097 | [-0.153 -0.041] | -3.458 | 50 | 0.001 |
| $\beta$ SoA on Confidence | 0.152 | [0.106 0.198] | 6.677 | 50 | 0.000 |
| $\beta$ disparity on SoA | 0.217 | [0.120 0.314] | 4.502 | 50 | 0.000 |
| $\beta$ disparity on RT | -0.136 | [-0.189 -0.082] | -5.089 | 50 | 0.000 |
| $\beta$ disparity on confidence | 0.151 | [0.089 0.213] | 4.887 | 50 | 0.000 |

| **Experiment 5** |  |  |  |  |  |
| --- | --- | --- | --- | --- | --- |
| **DV** | **M** | **C.I.** | **t** | **d.f.** | **p** |
| median SoA<sub>1</sub> | 1.182 | [-0.091 2.455] | 1.861 | 54 | 0.068 |
| $\beta$ dV<sub>1</sub> on RT | -0.222 | [-0.283 -0.161] | -7.307 | 54 | 0.000 |
| $\beta$ dV<sub>1</sub> on Consistency | 3.324 | [2.637 4.011] | 9.702 | 54 | 0.000 |
| $\beta$ dV<sub>1</sub> on Confidence | 0.355 | [0.296 0.415] | 11.937 | 54 | 0.000 |
| $\beta$ dV<sub>2</sub> on RT | -0.244 | [-0.307 -0.180] | -7.715 | 54 | 0.000 |
| $\beta$ dV<sub>2</sub> on Consistency | 4.010 | [3.213 4.807] | 10.086 | 54 | 0.000 |
| $\beta$ dV<sub>2</sub> on Confidence | 0.413 | [0.347 0.479] | 12.568 | 54 | 0.000 |
| $\beta$ dV<sub>3</sub> on RT | -0.282 | [-0.345 -0.219] | -8.963 | 54 | 0.000 |
| $\beta$ dV<sub>3</sub> on Consistency | 4.492 | [3.638 5.347] | 10.535 | 54 | 0.000 |
| $\beta$ dV<sub>3</sub> on Confidence | 0.479 | [0.420 0.538] | 16.292 | 54 | 0.000 |
| $\beta$ SoA<sub>1</sub> on RT | -0.141 | [-0.206 -0.076] | -4.351 | 54 | 0.000 |
| $\beta$ SoA<sub>2</sub> on RT | -0.122 | [-0.164 -0.081] | -5.942 | 54 | 0.000 |
| $\beta$ SoA<sub>1</sub> on Confidence | 0.284 | [0.217 0.350] | 8.573 | 54 | 0.000 |
| $\beta$ SoA<sub>2</sub> on Confidence | 0.163 | [0.109 0.216] | 6.052 | 54 | 0.000 |
| $\beta$ SoA on RT | -0.093 | [-0.154 -0.032] | -3.063 | 54 | 0.003 |
| $\beta$ SoA on Confidence | 0.213 | [0.154 0.272] | 7.219 | 54 | 0.000 |
| $\beta$ disparity on SoA | 0.197 | [0.110 0.283] | 4.557 | 54 | 0.000 |
| $\beta$ disparity on RT | -0.066 | [-0.132 0.000] | -2.003 | 54 | 0.050 |
| $\beta$ disparity on confidence | 0.144 | [0.071 0.217] | 3.964 | 54 | 0.000 |

---

**Figure S1.**
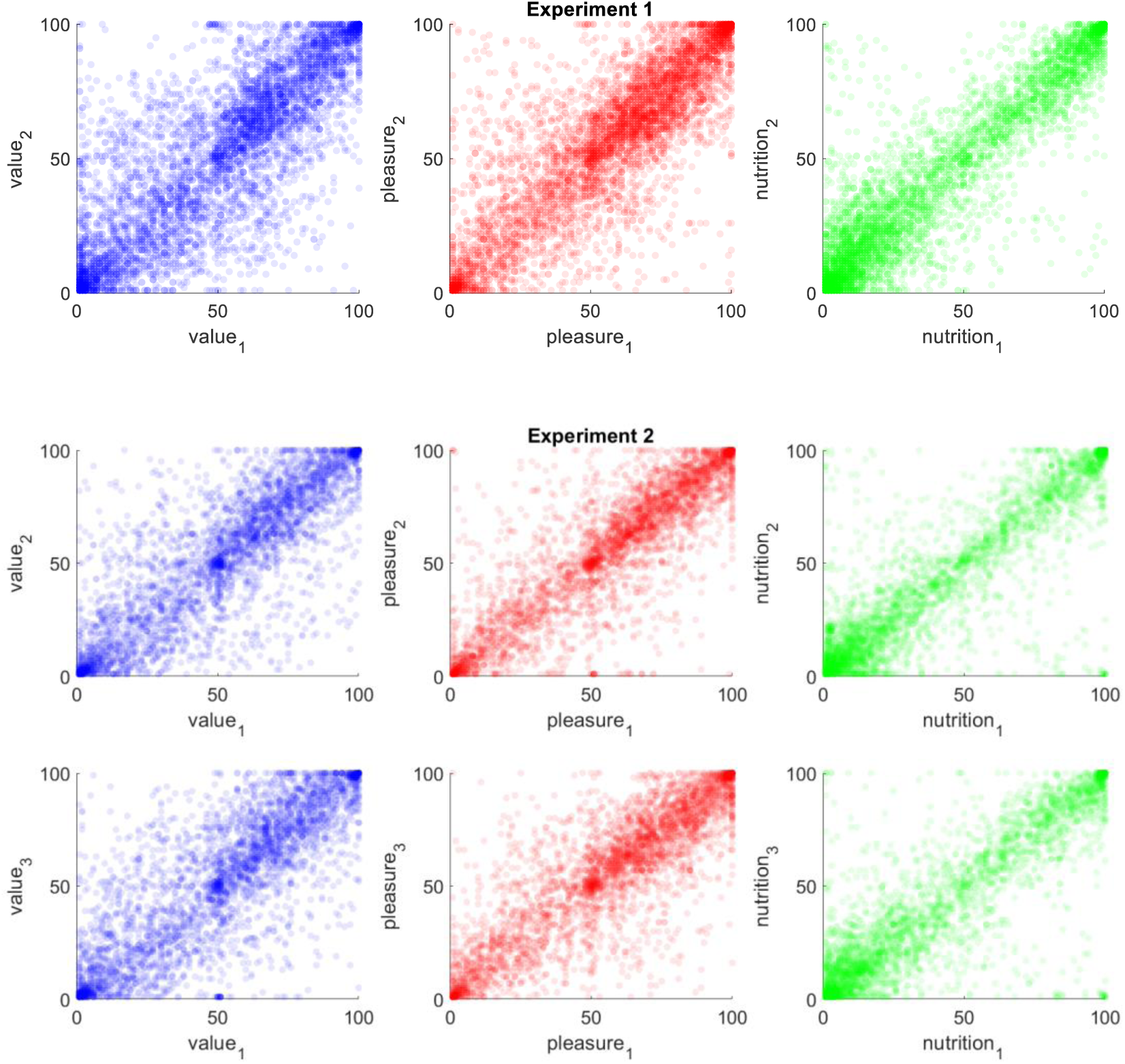
Test-retest reliability for ratings across experimental task sections. Left plots show overall value ratings (blue), middle plots show pleasure ratings (red), and right plots show nutrition ratings (green). The data for Experiments 3-5 were similar to the data for Experiment 2.

**Figure S2.**
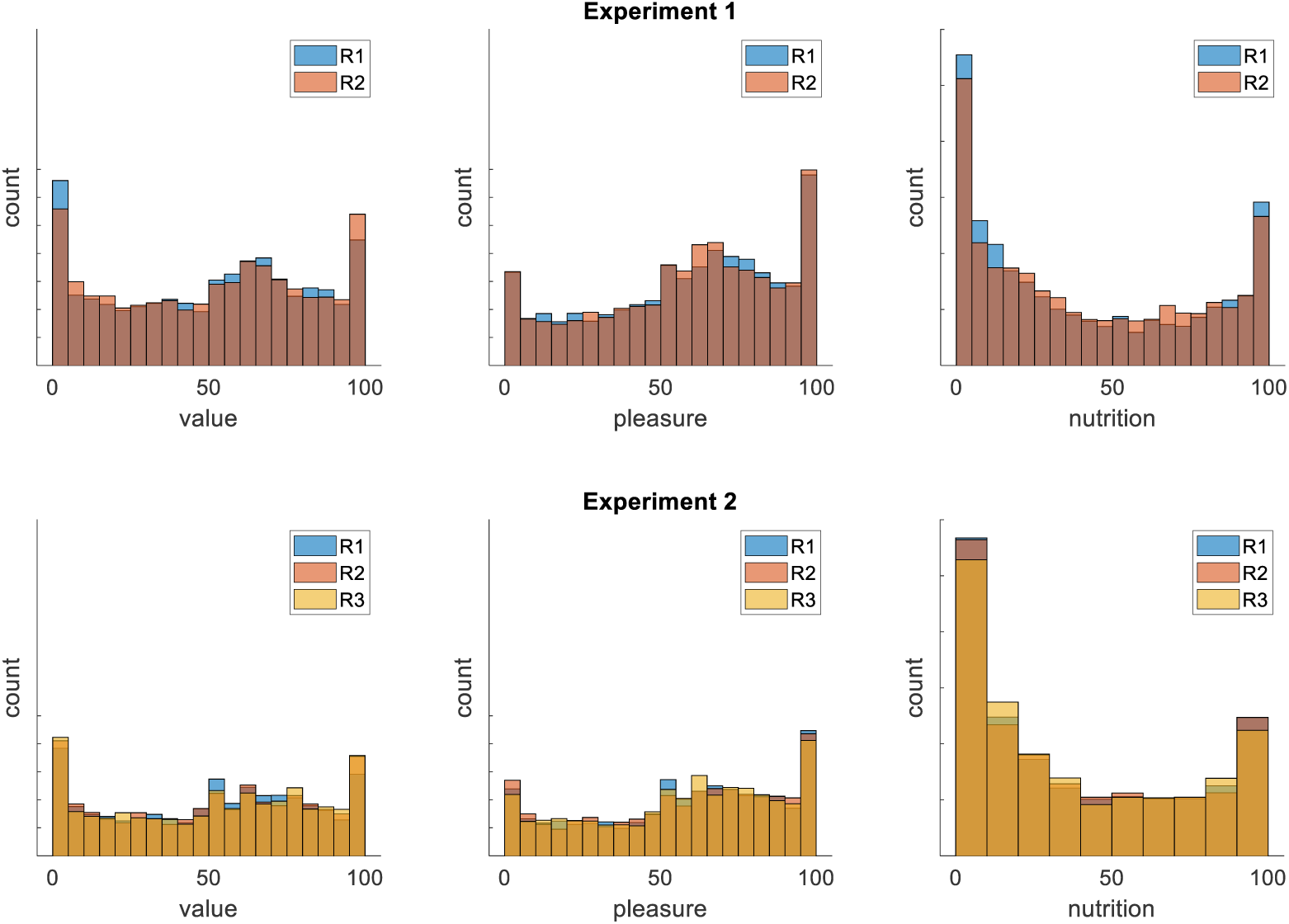
Histograms of ratings (overall value, pleasure, and nutrition) pooled across participants, separately for Experiment 1 (top row) and Experiment 2 (bottom row). R1 (blue) is for the first rating task, R2 (red) is for the second rating task, R3 (yellow) is for the third rating task. Notably, there is no difference in any of the distributions across tasks. The data for Experiments 3-5 were similar to the data for Experiment 2.

**Figure S3.**
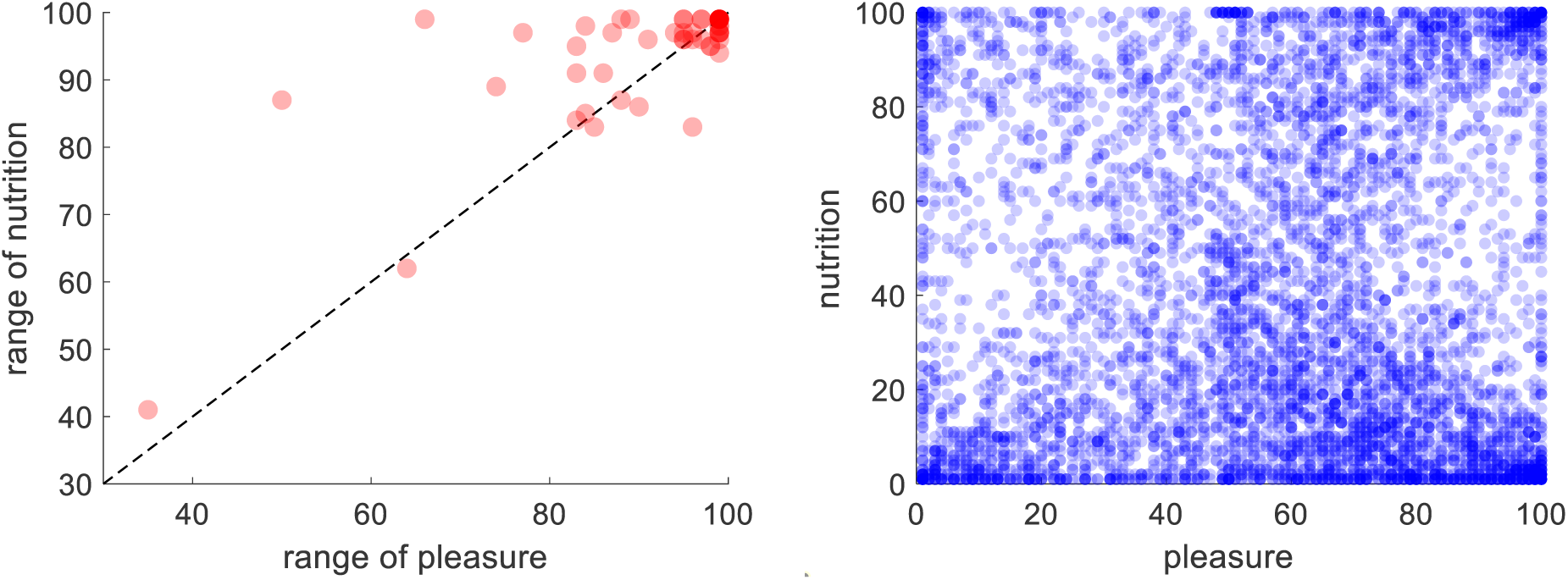
Across participants, pleasure and nutrition ratings occupied the full range of the scale. The left plot shows the range nutrition ratings against the range of pleasure ratings (each red dot indicates one participant). The right plot shows that the full pleasure-nutrition space was sufficiently covered, across participants (each blue dot represents one item). Here we show only the data for Experiment 1; the data for Experiments 2-5 were similar.

**Figure S4.**
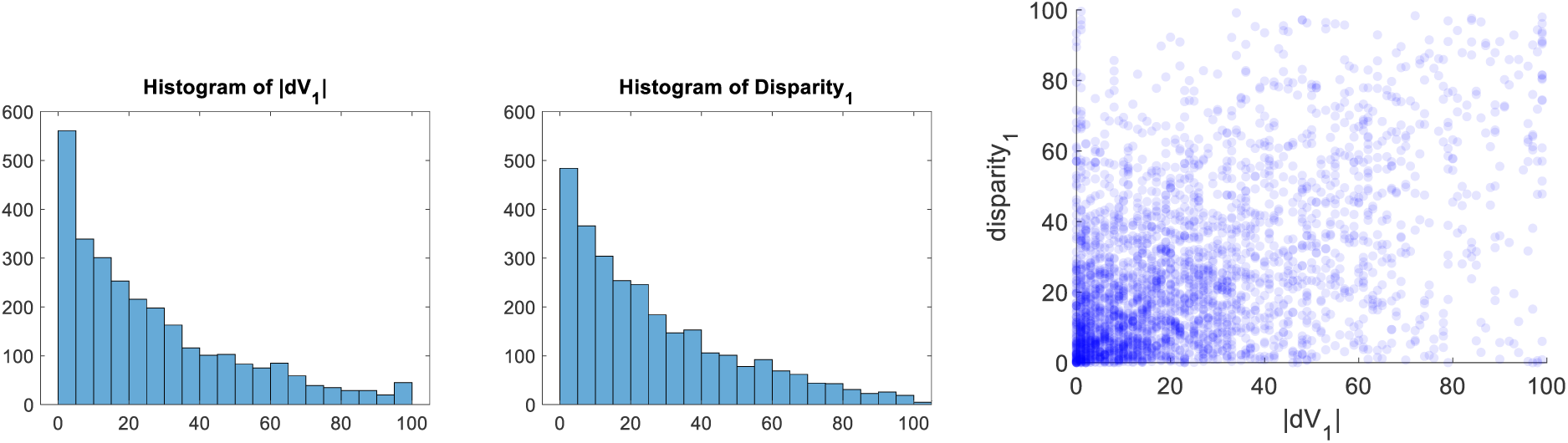
dV and disparity were similarly and sufficiently well distributed. Left and middle plots show histograms of dV and disparity (here shown for R1, across participants). Right plot shows that the full dV-disparity space was sufficiently well covered. Here we show only the data for Experiment 1; the data for Experiments 2-5 were similar.

**Figure S5.**
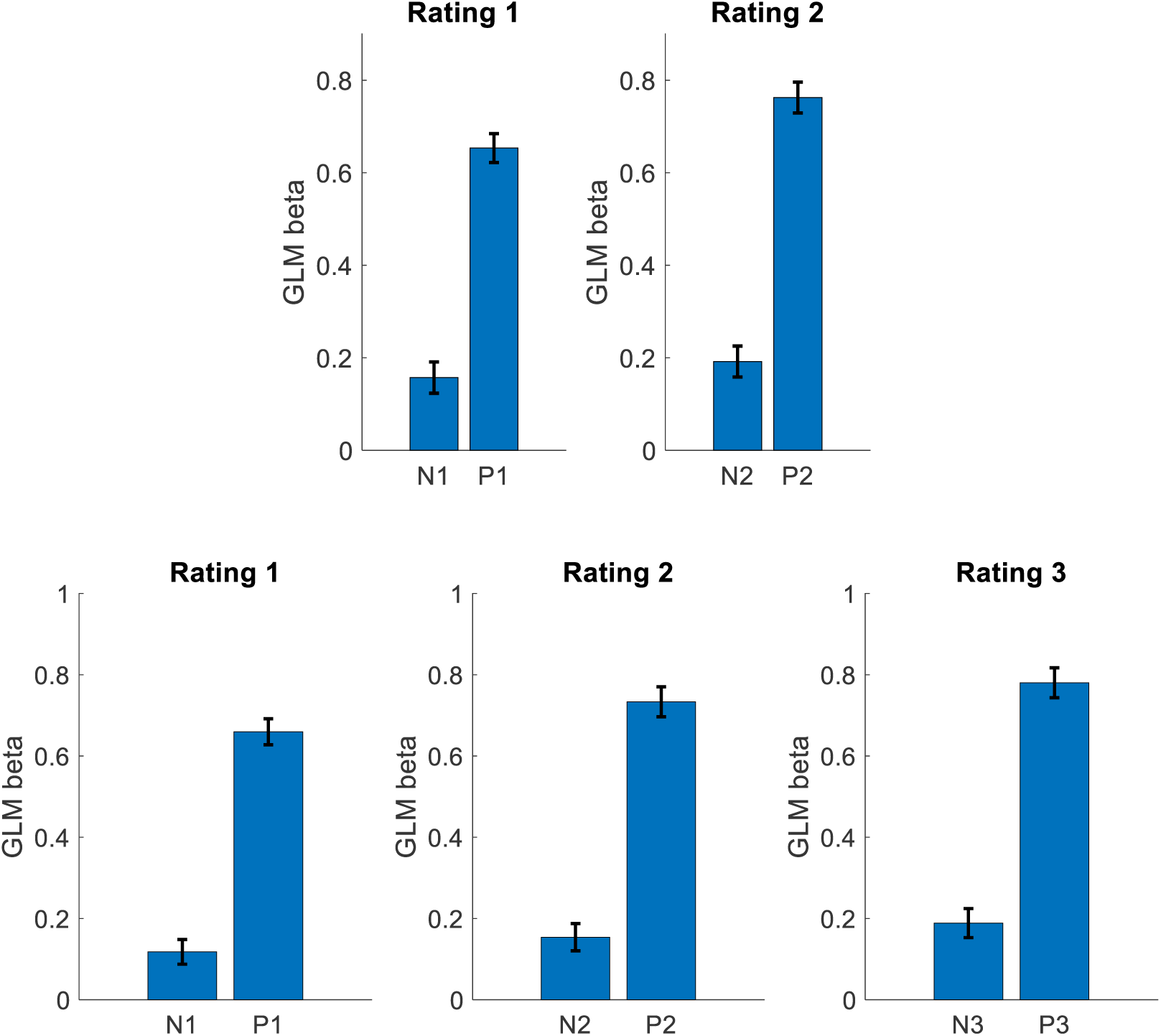
Importance weights of attributes contributing to overall value for Experiment 1 (top row) and Experiment 2 (bottom row), calculated using GLM regression of attribute ratings onto overall value ratings. The magnitude of the beta weights for both nutrition (N) and pleasure (P) increased from first to second rating task, and from second to third rating task, suggesting that deliberation (in the choice or similarity judgment tasks between each rating task) refined the subjective mapping from attribute values to overall value. The data for Experiments 3-5 were similar to the data for Experiment 2.

**Figure S6.**
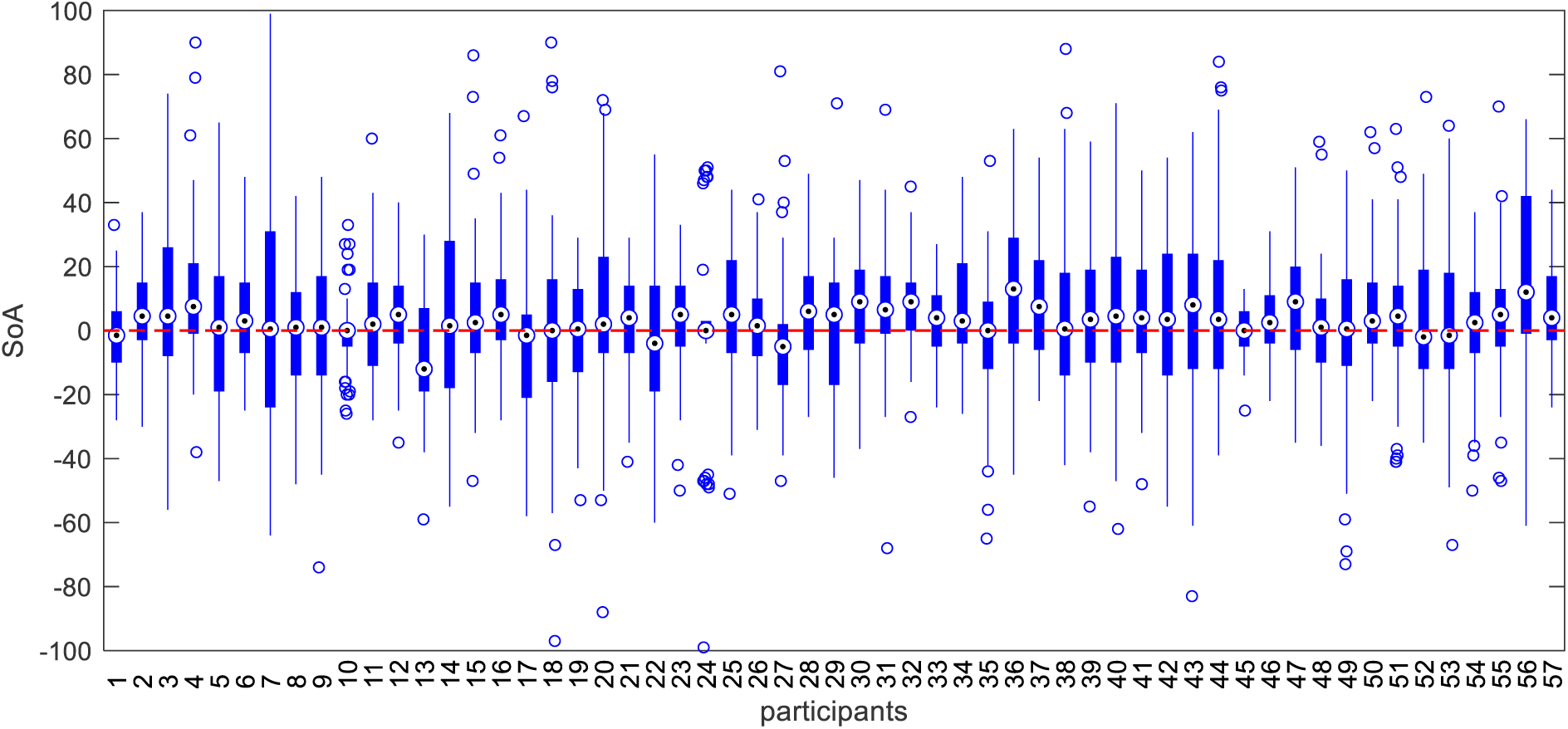
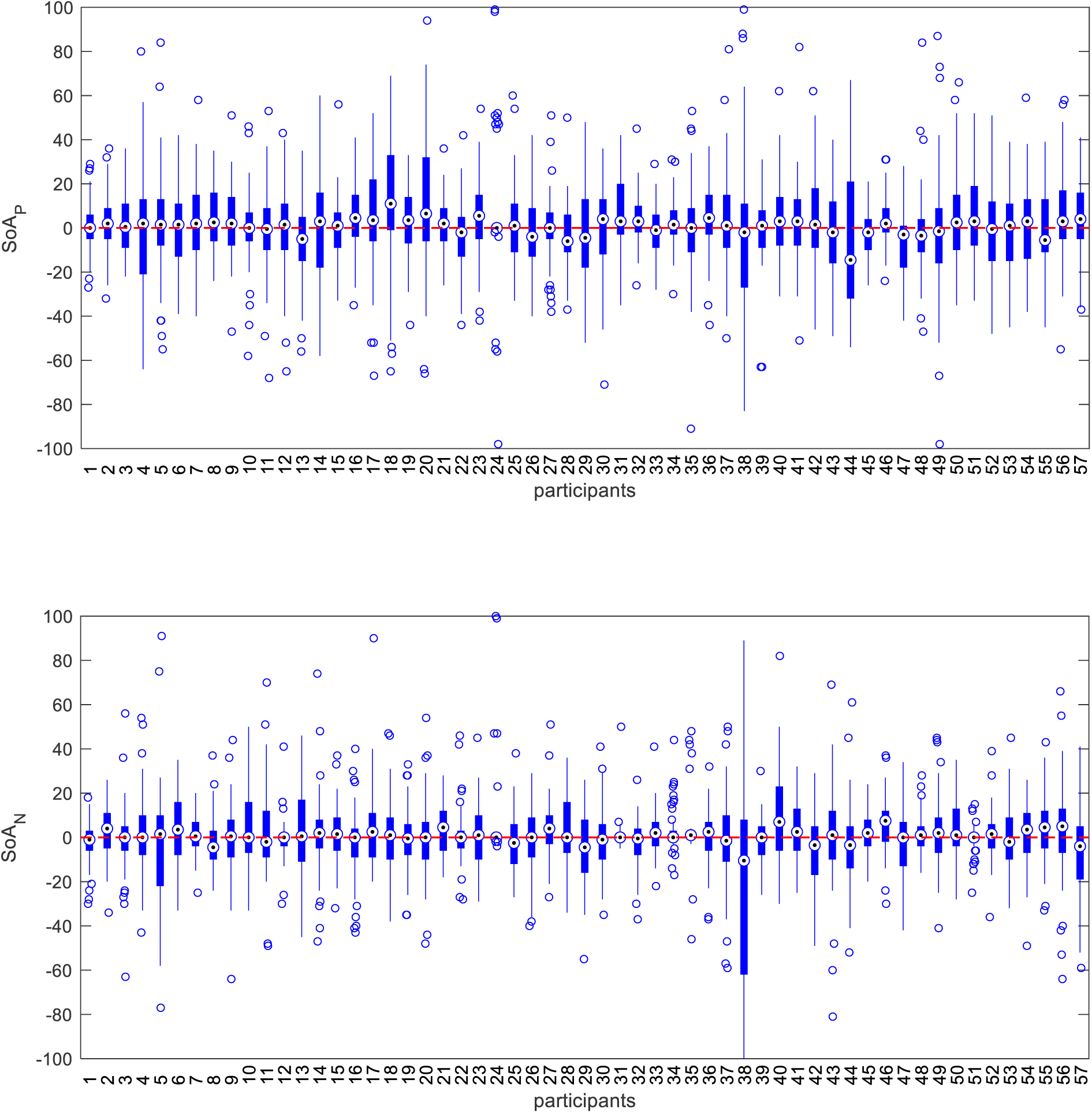
Within-participant distributions of spreading of alternatives for overall value (SoA, top plot), pleasure (SoA_P_, middle plot), and nutrition (SoA_N_, bottom plot). For each box, the central mark is the median, the edges of the box are the 25th and 75th percentiles, the whiskers extend to the most extreme data points the boxplot algorithm considers not to be outliers, and the outliers are plotted individually. Here we show only the data for Experiment 1; the data for Experiments 2-5 were similar.

**Figure S7.**
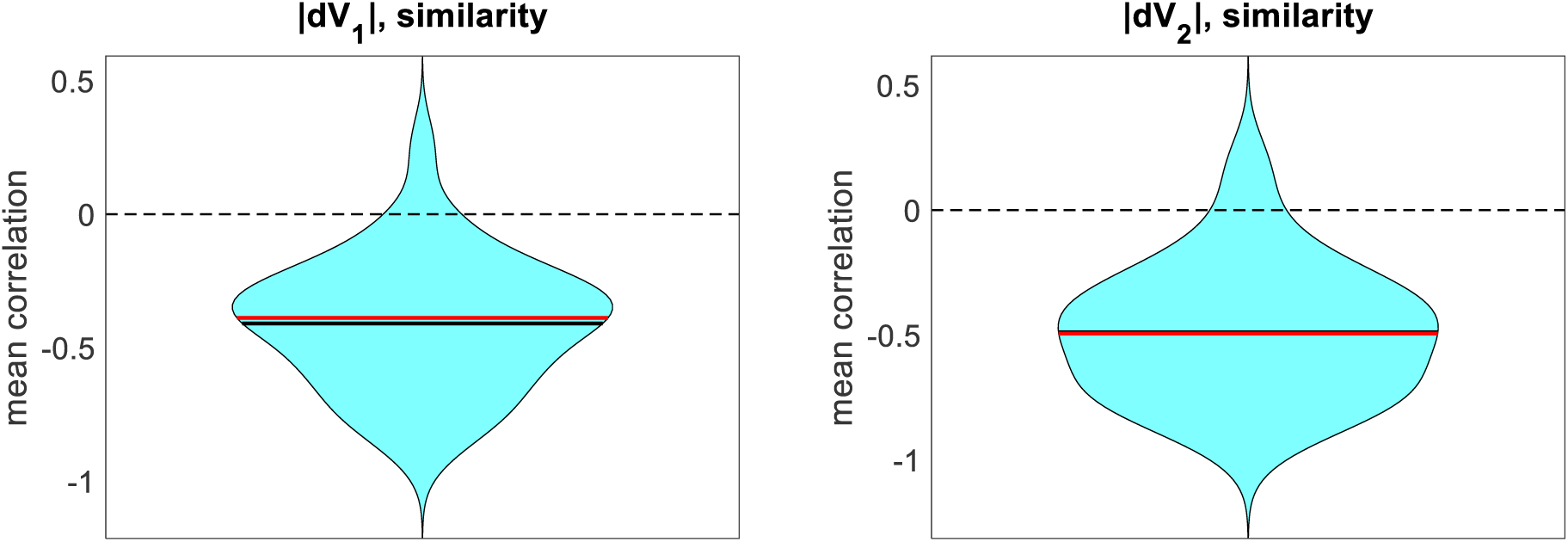
Similarity judgments in Experiment 2 capture value difference, as intended. Overall value difference for choice pair options (dV) correlated negatively with subjective reports of similarity for the same choice pair options. Left plot shows the correlation between similarity and pre-judgment ratings; right plot shows the correlation between similarity and post-judgment ratings.

## Footnotes

1 We do not include consistency here, because some choices would be classified as preference reversals (i.e., dV1 and dV3 would have opposite signs). These cannot necessarily be classified as errors, however, as the value refinements might have caused the decision makers to change their mind about their preferences on these trials. Thus, for these trials, choice consistency would be uncorrelated or anti-correlated with confidence and RT, although all three variables would be influenced by SoA.

## Notes

### Competing Interest Statement

The authors have declared no competing interest.

### Summary of Updates

several new experiments are now included, an important new finding is now discussed, detailed statistics are appended

